# A Calcium/Palmitoylation Switch Interfaces the Signaling Networks of Stress Response and Transition to Flowering

**DOI:** 10.1101/2021.08.05.455199

**Authors:** Hee Jin Park, Francisco Gamez-Arjona, Marika Lindahl, Rashid Aman, Irene Villalta, Raul Carranco, Chae Jin Lim, Elena García, Ray A. Bressan, Sang Yeol Lee, Clara Sánchez-Rodríguez, Jose M Pardo, Woe-Yeon Kim, Francisco J. Quintero, Dae-Jin Yun

## Abstract

The precise timing of flowering in adverse environments is critical for plants to secure reproductive success. We report a novel mechanism controlling the time of flowering by which the palmitoylation-dependent nuclear import of protein SOS3/CBL4, a Ca^2+^-signaling intermediary in the plant response to salinity, results in the selective stabilization of the flowering time regulator GIGANTEA inside the nucleus under salt stress, while degradation of GIGANTEA in the cytosol releases the protein kinase SOS2 to achieve salt tolerance. *S*-acylation of SOS3 was critical for its nuclear localization and the promotion of flowering, but dispensable for salt tolerance. SOS3 interacted with the photoperiodic flowering components GIGANTEA and FKF1 on the *CONSTANS* gene promoter to sustain the transcription of *CO* and *FT* under salinity. Thus, SOS3 acts as a Ca^2+^- and palmitoylation-dependent molecular switch that fine-tunes flowering in a saline environment through the shared spatial separation and selective stabilization of GIGANTEA. The SOS3 protein connects two signaling networks to co-regulate stress adaptation and time of flowering.

**Short summary:** S-acylation promoted the nuclear import of SOS3/CBL4 for the selective stabilization of the photoperiodic floral regulator GIGANTEA to fine-tune flowering time in a saline environment. Spatial separation of SOS3 acts as a molecular switch co-regulating stress adaptation and time of flowering.

## INTRODUCTION

Natural selection of different biological forms and functions occurs in the variable physical environments. Depending on the specific environment, different traits are favored for reproduction and perpetual survival of the species. For plants, extremes in the cardinal conditions such as light, temperature and most importantly, the quantity and quality of available water and nutrients are among the major drivers of natural selection (Maggio et al., 2018). Adaptive responses must be coupled to adjustments in the reproductive strategy to be favored by selection. Seasonal changes, especially in temperature and day length, provide key signals setting the time of flowering. However, depending on the dynamics of environmental stressors, transition to flowering is adjusted earlier or later to maximize the production of dormant structures (seeds) that can survive prolonged adverse episodes and eventually re-initiate a life cycle (Kazan and Lyons, 2016). Environmental stressors such as water and nutrient deprivation generally induce earlier flowering (Kazan and Lyons, 2016; Takeno, 2016), whereas salinity has been reported to delay flowering (Kim et al., 2007; Kim et al., 2013a; Li et al., 2007; Ryu et al., 2014).

In the model plant *Arabidopsis thaliana*, the major signaling systems that perceive environmental cues and initiate flowering converge on a few key integrators. *CONSTANS* (*CO*) is a central promoter of the photoperiodic flowering pathway through its enhancement of the expression of the floral-inductive *FLOWERING LOCUS T* (*FT*) in long-day conditions (Corbesier and Coupland, 2006; Turck et al., 2008). *CO* is transcriptionally regulated by the opposing action of activators and repressors controlled by the circadian clock, including GIGANTEA (GI) and CYCLING DOF FACTORS (CDF1, 2, 3 and 5), and post-transcriptionally by photoreceptors that affect CO protein stability (Fornara et al., 2009; Imaizumi et al., 2005; Sawa et al., 2007; Suarez-Lopez et al., 2001; Valverde et al., 2004). The abundance of CDF proteins is in turn depressed by the blue light receptor F-box E3 ubiquitin ligase FKF1 (FLAVIN-BINDING, KELCH REPEAT, F-BOX 1) (Fornara et al., 2009; Imaizumi et al., 2005; Suarez-Lopez et al., 2001). The clock protein GI interacts with and stabilizes FKF1 in a blue light-dependent manner, thus promoting the degradation of CDF proteins and *CO* expression in long days (Sawa et al., 2007). GI also forms a complex with and neutralizes *FT* repressors (Sawa and Kay, 2011) to enable *FT* transcription and promote transition to flowering (Mathieu et al., 2009).

Salt stress delays flowering time in Arabidopsis by repressing expression of *CO* and *FT* (Kim et al., 2007; Li et al., 2007). Moreover, the salt-induced BROTHER OF FT AND TBL1 (BFT) competes with FT for binding to FLOWERING LOCUS D (FD), a co-transcription factor of FT in flowering initiation, and contributes to late flowering (Ryu et al., 2014). In parallel, salinity promotes extension of vegetative growth by stabilizing DELLA proteins that act as repressors of cell proliferation and expansion, and of flowering (Achard et al., 2006). Other regulators mediating abiotic stress responses are also known to modulate flowering time and *vice versa*, but mechanistic insights are still largely missing (Park et al., 2016). Among these dual effectors is GI, that has emerged as a central hub coordinating the photoperiodic flowering pathway and stress responses against drought (Han et al., 2013; Riboni et al., 2013), cold (Cao et al., 2005; Fornara et al., 2015), salt (Kim et al., 2013a), light (Oliverio et al., 2007), and carbohydrate metabolism (Dalchau et al., 2011). The involvement of GI in stress responses includes transcriptional regulation of downstream genes (Fornara et al., 2015) and the interaction with circadian and other signaling components that in turn affect various physiological adaptations (Greenham and McClung, 2015; Park et al., 2016; Seo and Mas, 2015).

We have shown that GI controls salt tolerance through the direct association with key signaling components of the salinity stress response (Kim et al., 2013a; Park et al., 2016). In response to high salinity, plants utilize the SOS (Salt Overly Sensitive) pathway to maintain ion homeostasis. The core components of the SOS pathway comprise the Na^+^/H^+^ antiporter SOS1, the Ser/Thr protein kinase SOS2/CIPK24, and two alternative calcium binding proteins, SOS3/CBL4 and SCaBP8/CBL10, that activate and recruit SOS2 to cellular membranes (Qiu et al., 2002; Quintero et al., 2002; Quan et al., 2007; Quintero et al., 2011). SOS2 activates Na^+^ efflux by phosphorylating SOS1 at its C-terminal autoinhibitory domain (Quintero et al., 2002; Quintero et al., 2011). In regular growth conditions, GI makes a complex with and inhibits SOS2 (Kim et al., 2013a). Salt stress causes the degradation of GI protein by the 26S proteasome and the release of SOS2, which is then free to interact with SOS3, activate SOS1 and mount a successful adaptation to the saline environment. The removal of GI leads to exceptional salt tolerance at least in part by mimicking Na^+^-induced GI degradation, whereas plants overexpressing *GI* exhibit a salt-sensitive phenotype by sequestering SOS2. The precise mechanism triggering the dissociation of the GI-SOS2 complex and GI degradation under salt stress has not been resolved, although indirect evidence suggested that the Ca^2+^-sensor protein SOS3 played a role since excess SOS3 interfered with GI-SOS2 complex formation (Kim et al., 2013a). Moreover, SOS3 has been reported to have an indeterminate role in influencing flowering time as the *sos3-1* mutant, which has impaired calcium binding, showed late flowering under salt stress (Ishitani et al., 2000; Li et al., 2007). The molecular basis of this phenotype has remained unexplained.

We have addressed the molecular mechanism by which SOS3 helps resetting the flowering time under salt stress. We show here that SOS3 acts as a crucial regulator of flowering under saline stress through a mechanism that involves the stabilization of GI specifically inside the nucleus. Under normal growth conditions, GI partitions between the nucleoplasm and cytoplasm. Upon salinity stress, only cytoplasmic GI is degraded, thereby releasing SOS2 to mount the salt stress response, whereas nuclear GI remains stable in physical association with SOS3, eventually leading to flowering. Notably, S-acylation with fatty acids, commonly known as protein palmitoylation (Hemsley, 2020), of SOS3 is required for the nuclear import of SOS3 but dispensable for the interaction with GI. We also demonstrate the participation of SOS3 in the GI-FKF1 transcriptional complex that promotes transcription of *CO*, a crucial floral activator. These results reveal the molecular linkages of networks controlling salinity stress responses and the adaptive initiation of flowering under adverse environments. They also reveal a novel mechanism for transcriptional regulation of flowering determinants by a Ca^2+^-activated protein whose nuclear import is controlled by S-acylation.

## RESULTS

### SOS3 controls flowering under saline stress through the *CO/FT* pathway

Previously we have shown that GI, which promotes photoperiodic-dependent flowering in long days (LDs), also functions to restrain the activity of the SOS pathway by sequestering SOS2 (Kim et al., 2013a). Upon salt stress, the GI-SOS2 complex dissociates and free GI degrades to delay flowering. This creates a reciprocating on/off mechanism coordinating signal networks of stress response and time of flowering. Under regular growth conditions, *sos1-1*, *sos2-2* and *sos3-1* mutants flowered as the wild-type (Kim et al., 2013a; Li et al., 2007). However, the flowering of the *sos3-1* mutant was delayed under salt stress compared to wild-type, *sos1-1* and *sos2-2* (Figure 1). The *sos1-1* mutant exhibited maximal sensitivity to 30 mM NaCl among all genotypes tested but still flowered normally, indicating that delayed flowering in *sos3-1* plants was not a consequence of Na^+^ toxicity.

**Figure 1.**
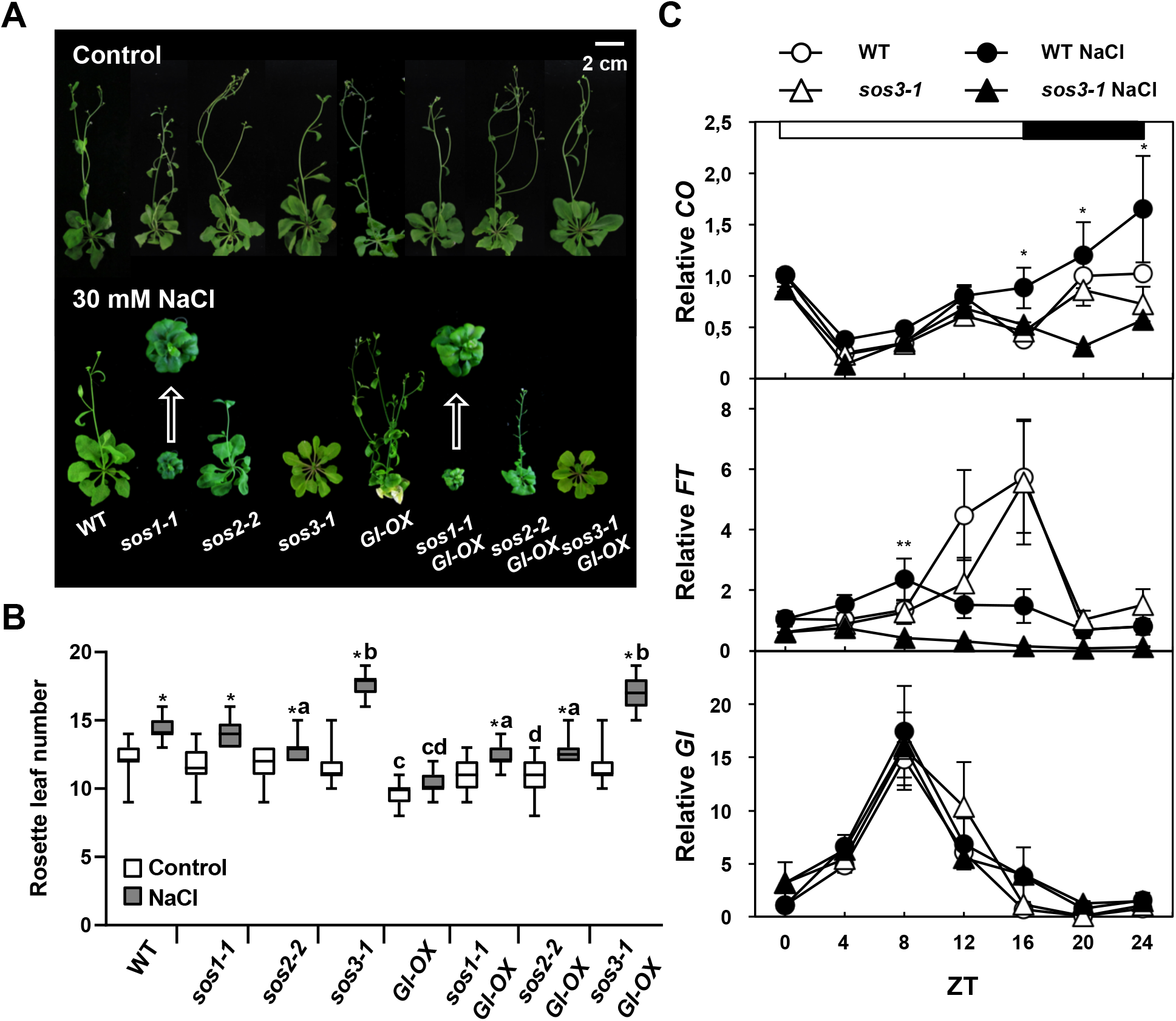
SOS3 controls flowering under salt stress through the CO/FT pathway. (**A**) Effect of salt on the flowering time in wild-type Col-*gl1* (WT), and mutants *sos1-1*, *sos2-2* and *sos3-1* overexpressing or not *GIGANTEA (GI-OX*). Eight-day old seedlings were transferred to MS media supplemented or not with 30 mM NaCl. The photographs were taken after bolting. Representative plants are shown. (**B**) Rosette leaf number at bolting time of plants grown with and without salt, as in (A) to score flowering time. Data is shown as box plots: center lines show the medians; box limits indicate the 25th and 75th percentiles; whiskers extend to the minimum and maximum (n=16). Asterisks indicate significantly different means of samples with and without salt for each genotype, and letters indicate differences of mutant and transformed lines with the corresponding sample of the WT with and without salt at p<0.01, Fisher’s LSD test; means with the same letter are statistically similar. (**C**) Transcript levels of *CO*, *FT* and *GI* in wild-type (Col-gl1) and *sos3-1* mutant. Two-week old plants grown in long-days were left untreated (open symbols) or treated with 100 mM NaCl (filled symbols) at the beginning of the light period (ZT0), and harvested every 4 h. Total RNA was isolated and transcript levels of *CO*, *FT* and *GI* were measured by qRT-PCR and normalized to that of *At5g12240*. Errors bars represent means ± SEM from at three replicates with three technical replicates each. Asterisks indicate significant differences between genotypes with the same treatment, * p<0.05, ** p<0.01, by Fisher’s LSD test. The white and black bars on top indicate light and dark periods.

Ultimately, salt stress delays flowering because of reduced transcript levels of the GI-regulated floral activator *FT* (Li et al., 2007; Kim et al., 2013a; Sawa et al., 2007; Sawa and Kay, 2011). In the wild-type, salt stress altered the photoperiodic oscillation of *CO* transcripts and instead promoted the increase of *CO* throughout dusk and night (Figure 1C). *FT* levels followed the opposite trend, with a marked decline after midday (ZT8) and losing the maxima at dusk (ZT16) typical of untreated controls (Cerdan and Chory, 2003; Suarez-Lopez et al., 2001). *CO* and *FT* transcripts in the *sos3-1* mutant followed wild-type dynamics under control conditions, but salt treatment reduced *CO* at night, thus departing from the wild-type behavior, and abated further the *FT* transcript levels compared to the wild-type (Figure 1C). The diurnal dynamics of the *GI* transcript was not affected by salt or the *sos3-1* mutation (Figure 1C). Salinity did not alter the transcript levels of other flowering genes such as *FKF1*, *SOC1*, *FLD* (*FLOWERING LOCUS D*), *FLC*, and *FCA* (*FLOWERING TIME CONTROL PROTEIN FCA*) (Supplemental Figure S1) (Li et al., 2007). These results indicate that SOS3 not only mediates adaption to salinity through the SOS pathway but also participates in resetting flowering time through the CO-FT module under salt stress.

### SOS3 stabilizes GI in the nucleus under salt stress

Accumulation of the GI protein in late afternoon of LDs promotes the transcription of floral activators *CO* and *FT* (Sawa et al., 2007; Sawa and Kay, 2011), and *GI* overexpression leads to early flowering (David et al., 2006). Overexpression of *GI* in *sos1-1* and *sos2-2* mutant backgrounds promoted unconditional early flowering but failed to suppress the delayed flowering of the *sos3-1* mutant under salt stress (Figure 1A and 1B). This suggests that promotion of flowering during salt stress by GI strictly requires a functional SOS3.

GI is a nucleo-cytoplasmic protein, and forced spatial segregation of GI into nuclear or cytosolic compartments results in different outputs of GI function (Kim et al., 2013b). Transgenic plants exclusively expressing a recombinant GI protein fused to a nuclear localization signal (*GIpro:GI-GFP-NLS* in *gi-2* mutant, henceforth *GI-NLS*) resulted in unconditional early flowering compared to wild-type or control transgenic plants expressing nucleo-cytoplasmic *GIpro:GI-GFP* (Kim et al., 2013b). Conversely, transgenic plants expressing a preferentially cytoplasmic GI protein fused to a nuclear export signal (*GIpro:GI-GFP-NES* in *gi-2*, henceforth *GI-NES*) exhibited late flowering due to nuclear exclusion of GI. This late flowering of GI-NES plants was exacerbated under salt stress and resembled that of the untransformed *GI*-deficient mutant *gi-2* (Figure 2A and 2B). These results indicate that only the nuclear GI pool appears to be in control of promoting the photoperiodic flowering pathway, and that the salt-induced delay in flowering known to result from a decrease in the steady-state levels of GI protein (Kim et al., 2013a) affects primarily the cytosolic GI pool. Accordingly, we found that salt-induced degradation of a tagged GI-HA protein occurred only in the cytosol and not in the nucleus (Figure 2C and Supplemental Figure S2A). This observation was confirmed with salt-treated tobacco leaves that transiently expressed *GI-GFP* (Supplemental Figure S2B). Collectively, these results indicate that import to and preservation of GI stability inside the nucleus is critical to ensure flowering under salinity stress.

**Figure 2.**
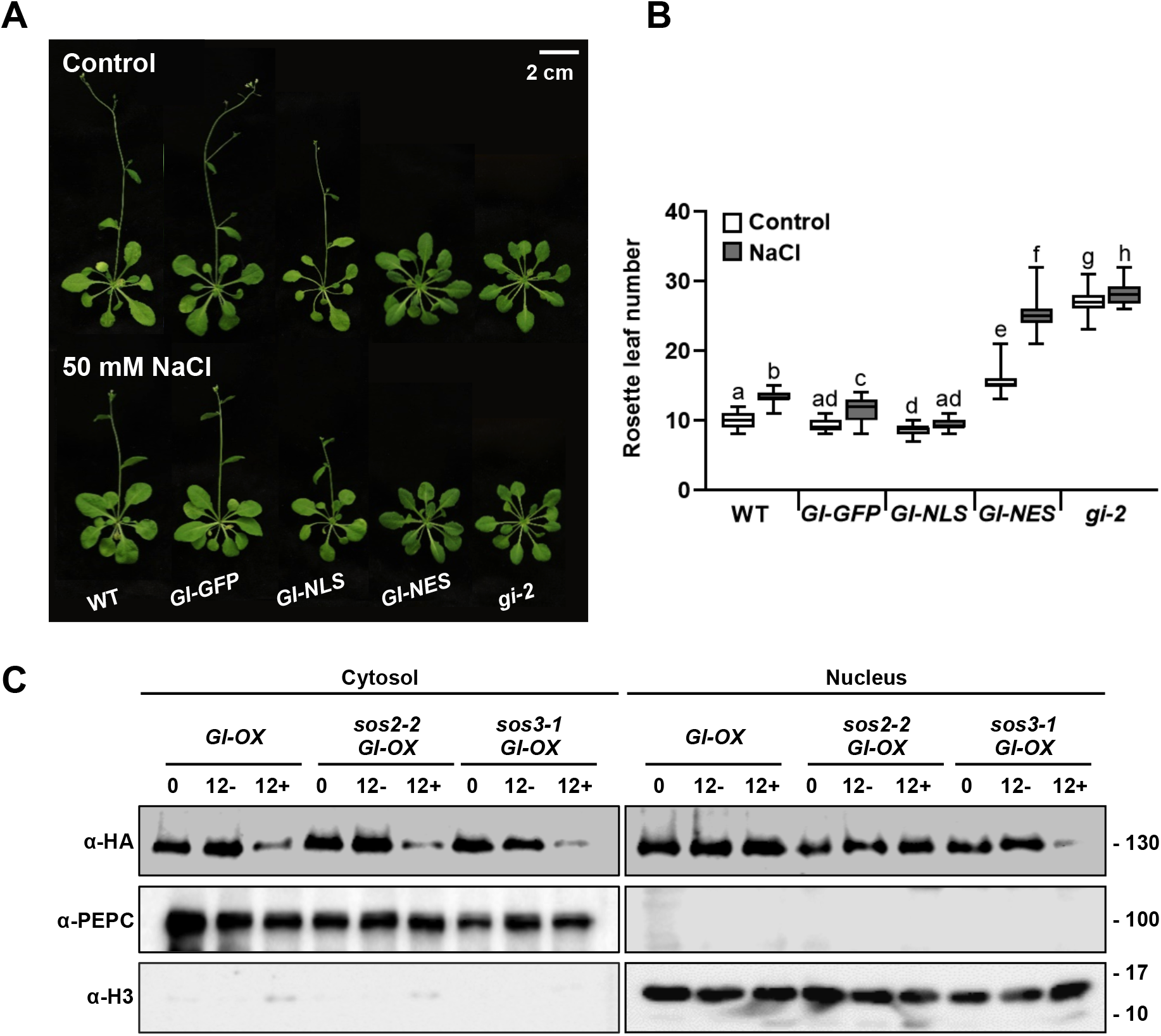
Stability of the nuclear fraction of GI controls time of flowering. (**A**) Eight-day old plants of wild-type Col-0 (WT) and the *gi-2* mutant transformed with *GI-GFP, GI-NLS and GI-NES* were grown in LDs treated or not with 50 mM NaCl. (**B**) Flowering time was counted as the rosette leaf number at bolting. Shown are the Box plots: center lines show the medians; box limits indicate the 25th and 75th percentiles; whiskers extend to the minimum and maximum (n≥15). Letters indicate significantly different means at p<0.01, by Fisher’s LSD test. (C) Cytosolic and nuclear proteins were extracted from 2-week old *GI-OX*, *sos2-2 GI-OX* and *sos3-1 GI-OX* plants treated with (indicated as 12+) or without (indicated as 12-) 100 mM NaCl for 12 h. Immunoblots with HA antibody were performed to detect GI protein. α-PEPC and α-H3 antibodies were used for cytosolic and nuclear markers, respectively. This experiment was repeated three times with similar results.

To examine whether SOS3 is involved in the salt-regulated GI stability, *GI-OX*, *sos2-2 GI-OX* and *sos3-1 GI-OX* transgenic plants were treated with 100 mM NaCl for 12 h starting at ZT2, and cytosolic and nuclear proteins were extracted. Salt-induced degradation of cytosolic GI was found in all plant lines (Figure 2C). By contrast, reduction of the nuclear GI pool was found only in *sos3-1 GI-OX* transgenic plants. This result suggests that SOS3 is needed for the stabilization of the GI protein within the nucleus under salt stress. We also tested whether CBL10/SCaBP8 (CALCINEURIN B-LIKE10/SOS3-LIKE CALCIUM BINDING PROTEIN8), a homolog of SOS3/CBL4 that interacts with SOS2 to impart salt tolerance (Quan et al., 2007) is involved in salinity-delayed flowering. Unlike *sos3-1*, the salt-induced delay in flowering was not observed in the *cbl10* mutant (Supplemental Figure S3).

### SOS3 interacts with GI in a calcium-dependent manner

Next, we tested whether salinity influenced the interaction of SOS3 with GI. Co-immunoprecipitation (co-IP) from tobacco leaves showed that GI-HA interacted with SOS3-MYC. The interaction was enhanced by 100 mM NaCl or 3 mM Ca^2+^ treatments, whereas EGTA suppressed the interaction (Figure 3A and 3B). Moreover, the mutant protein SOS3-1 bearing a three-amino acid deletion in the third EF-hand motif that abrogates interaction with SOS2 (Guo et al., 2004), also failed to interact with GI (Figure 3C). This suggests that Ca^2+^ promotes the interaction of SOS3 and GI. The Ca^2+^-dependent interaction of SOS3 with GI was confirmed by BiFC in tobacco (Figure 3D). The number of fluorescent nuclei and total fluorescence per area unit were counted as indicators of interaction strength (Figure 3E and Supplemental Figure S4A). Both NaCl and Ca^2+^ enhanced the interaction of GI and SOS3, whereas EGTA repressed the interaction. Again, GI did not interact with the mutant protein SOS3-1 (Figure 3D and 3E; control of protein expression in Supplemental Figure S4B), indicating that Ca^2+^ binding of SOS3 is important for interaction with GI.

**Figure 3.**
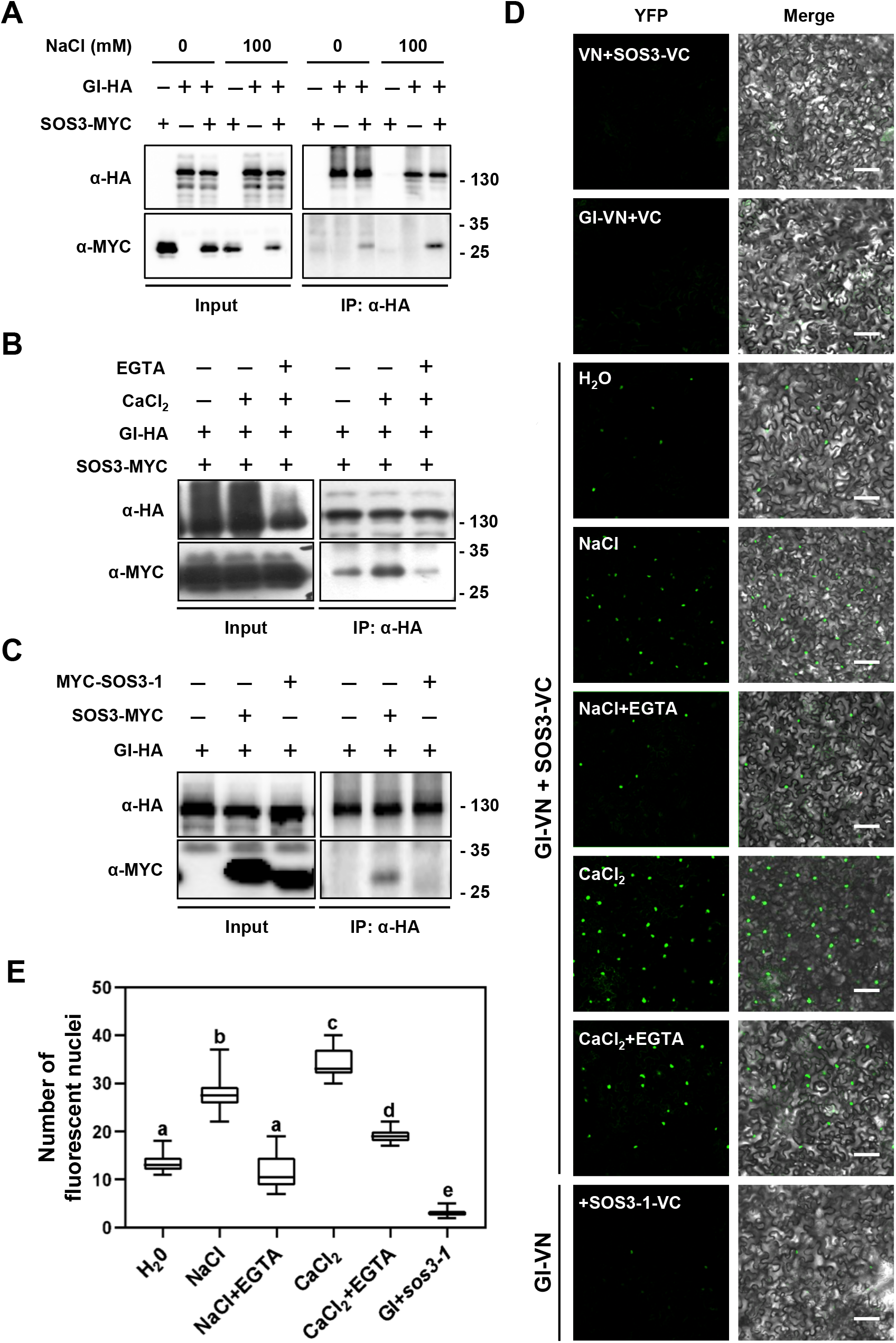
Salt- and Ca^2+^-dependent interaction of SOS3 with GI. (**A**) Co-immunoprecipitation of SOS3 and GI. Tobacco leaves transiently expressing SOS3-MYC and GI-HA were treated with 100 mM NaCl for 8 h and total proteins were pulled down with HA antibodies (α-HA). The SOS3 protein was detected by MYC antibodies (α-MYC). (**B**) Ca^2+^ effect on the interaction between SOS3 and GI. Tobacco leaves transiently expressing SOS3-MYC and GI-HA were treated with 3 mM CaCl_2_, or with 3 mM CaCl_2_ and 2 mM EGTA. (**C**) The SOS3-1 protein with a mutated EF-hand motif cannot bind to GI. (**D**) BiFC of GI and SOS3. Tagged GI-VN and SOS3-VC were transiently expressed in tobacco leaves and plants were treated for 6 to 8 h with 100 mM NaCl or 3 mM CaCl_2_ with or without 2 mM EGTA. Fluorescent signals were detected under confocal laser scanning microscope. Bar represents 100 µm. (E) The number of fluorescent nuclei in five images (0.4 mm^2^) of three biological replicas was counted and the results are shown as box plots: center lines show the medians; box limits indicate the 25th and 75th percentiles; whiskers extend to the minimum and maximum (n≥10). Letters indicate significantly different means, *p* <0.001 by Fisher’s LSD test, (n≥10); means with the same letter are statistically similar.

### S-acylation of SOS3 is crucial for nuclear import to ensure flowering under salt stress

Like GI, SOS3 is a nucleo-cytoplasmic protein (Batistič et al., 2010). N-myristoylation of SOS3 at Gly-2 is essential for the function of SOS3 in salt tolerance (Ishitani et al., 2000; Quintero et al., 2002). In addition, SOS3 has been suggested to undergo S-acylation at residue Cys-3 (Held et al., 2011). Therefore, we first confirmed that SOS3 is S-acylated *in vivo* and then tested whether N-myristoylation and S-acylation of SOS3 influenced protein localization and salt-induced delay of flowering. S-acylation at Cys-3 of wild-type SOS3 and mutant proteins G2A, C3A and G2A/C3A expressed in tobacco was tested by the acyl resin-assisted capture (acyl-RAC) method (Chai et al., 2019). Free cysteines in proteins were blocked with N-ethylmaleimide (NEM) prior to treatment or not with hydroxylamine (HyA), which breaks cysteine thioester bonds with fatty acids, and then proteins were attached covalently to the resin matrix through the newly formed cysteine thiols. Proteins were considered to be S-acylated if retention was observed only upon HyA treatment. Results demonstrated that SOS3 was S-acylated at Cys-3 and that this modification took place independently of myristoylation of Gly-2 (Figure 4A). The lower recovery of SOS3-G2A compared to SOS3 might due to decreased accessibility of the non-myristoylated proteins to PATs, which are integral membrane proteins (Rana et al., 2018). Since protein S-acylation is highly conserved in eukaryotes, SOS3 proteins (WT, G2A, C3A and G2A/C3A) were also recovered from yeast cells and the presence of *S*-linked fatty acids were analyzed by blocking thiol groups in SOS3 proteins with NEM, then treating with HyA, and finally cross-linking methyl-(PEG)_24_-maleimide to re-exposed cysteine thiols. A band shift was observed only in SOS3 and SOS3-G2A proteins, but not in proteins bearing the C3A mutation (Figure 4B). This result recapitulated the S-acylation pattern at Cys-3 of SOS3 proteins found in plants (Figure 4A).

**Figure 4.**
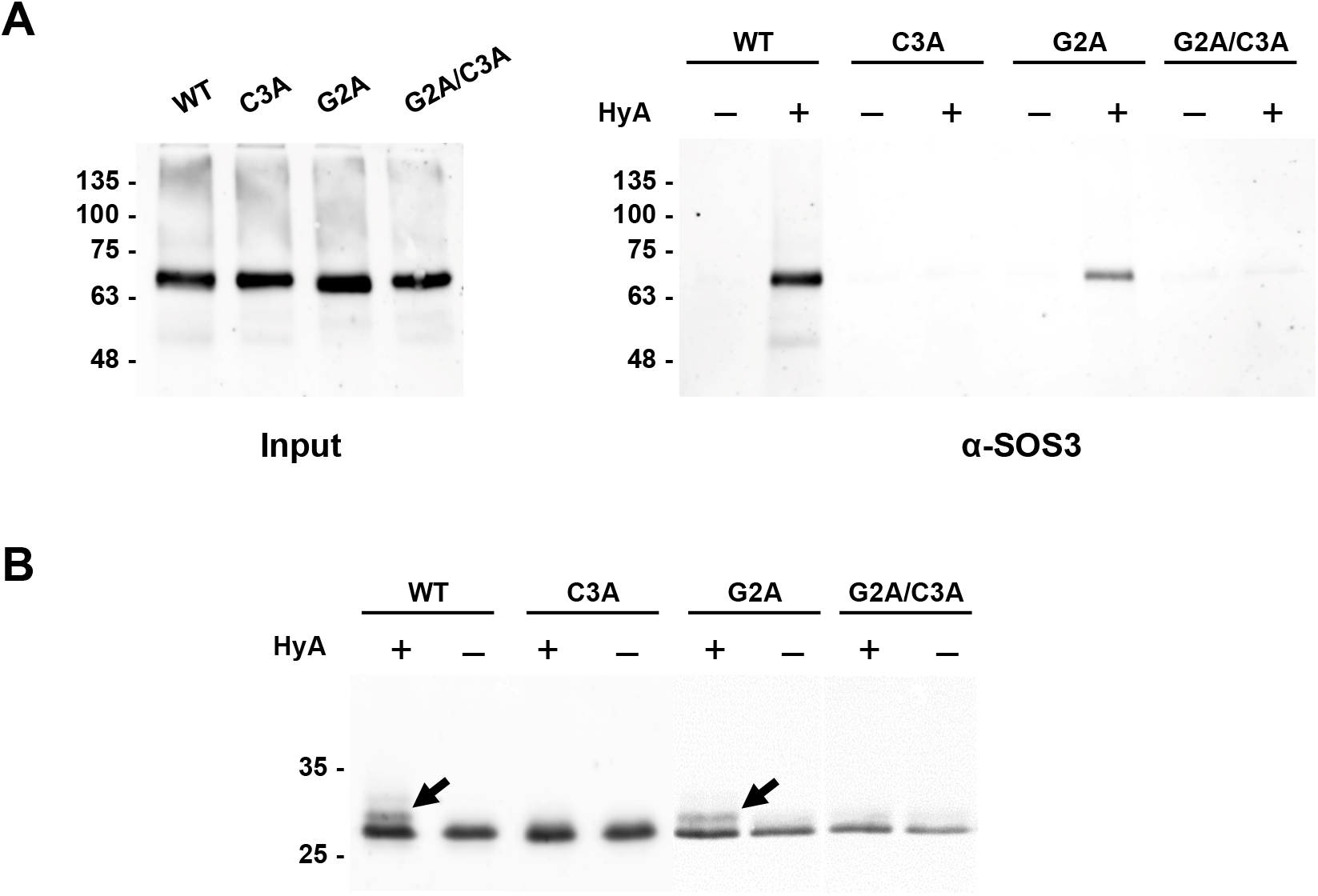
*S*-acylation of SOS3 at Cys-3. (**A**) Wild-type SOS3 and mutant variants G2A, C3A and the double mutant G2A/C3A, were expressed transiently in *N. benthamiana*. Leaf extracts were treated with 30 mM N-ethylmaleimide (NEM) under denaturing conditions to block free cysteine thiols and proteins were acetone precipitated. Resuspended proteins were incubated with thiopropyl-sepharose 6B in the presence (+) or absence (-) of 0.5 M hydroxylamine (HyA) to break palmitoyl thioester bonds. Right, covalently bound proteins were eluted with 50 mM DTT and probed by western blot using α-SOS3 antibody. Left, control of total leaf proteins applied as input for acyl-RAC, probed with α-SOS3 antibodies. Each lane contains proteins corresponding to 0.5 mg leaf tissue. (**B**) Wild-type SOS3 and mutant proteins G2A, C3A and G2A/C3A were expressed in yeast. Protein extracts were treated with NEM to block free cysteine thiols, and thereafter incubated in the presence (+) or absence (–) of hydroxylamine (HyA), precipitated with TCA and resuspended in 10 mM methyl-PEG24-maleimide, which alkylates newly formed cysteine thiols. Proteins resolved in SDS-PAGE were subjected to western blot analysis using the α-SOS3 antibody. The theoretical mass of SOS3 with 6xHis tag is 26.5 kDa and MM(PEG)_24_ adds 1.24 kDa for each cysteine alkylated (arrowheads).

We next generated transgenic plants expressing *35S:SOS3* (*SOS3-OX*), *35S:SOS3-G2A* (*SOS3-G2A*, no myristoylation), and *35S:SOS3-C3A* (*SOS3-C3A*, no S-acylation) in the *sos3-1* mutant, and tested their flowering time. All plants flowered at similar time in control conditions. Upon salt treatment, plants with constructs *SOS3-OX* and *SOS3-G2A* complemented the salt-induced flowering delay specific of *sos3-1* and had a flowering time similar to wild-type (Figure 5A and 5B), but expression of *SOS3-C3A* could not suppress this trait. Hence, we checked whether myristoylation and S-acylation of SOS3 were important for the interaction with GI. Total proteins extracted from tobacco leaves transiently expressing *SOS3-GFP*, *SOS3-G2A-GFP* or *SOS3-C3A-GFP* together with *GI-HA* were used for co-IP. Both SOS3-G2A-GFP and SOS3-C3A-GFP were able to interact with GI protein similarly to SOS3-GFP (Figure 5C). Notably, when expressed in *N. benthamiana* leaves, SOS3 and SOS3-G2A displayed a nucleo-cytoplasmic distribution but the nuclear import of SOS3-C3A mutant was suppressed (Figure 6A-B). The nuclear interaction of GI with SOS3-C3A was also severely reduced, although S-acylation of SOS3 was not strictly required for SOS3-GI interaction in co-IP and BiFC assays (Figure 5C-D, and Supplemental Figure S5). The GI complex with SOS3-C3A localized in a perinuclear rim suggesting aborted nuclear import and retention of the complex in the perinuclear ER. Together, these results evidence that S-acylation of SOS3 directs nuclear import of the SOS3-GI complex, which is required for ensuring flowering under salt stress conditions.

**Figure 5.**
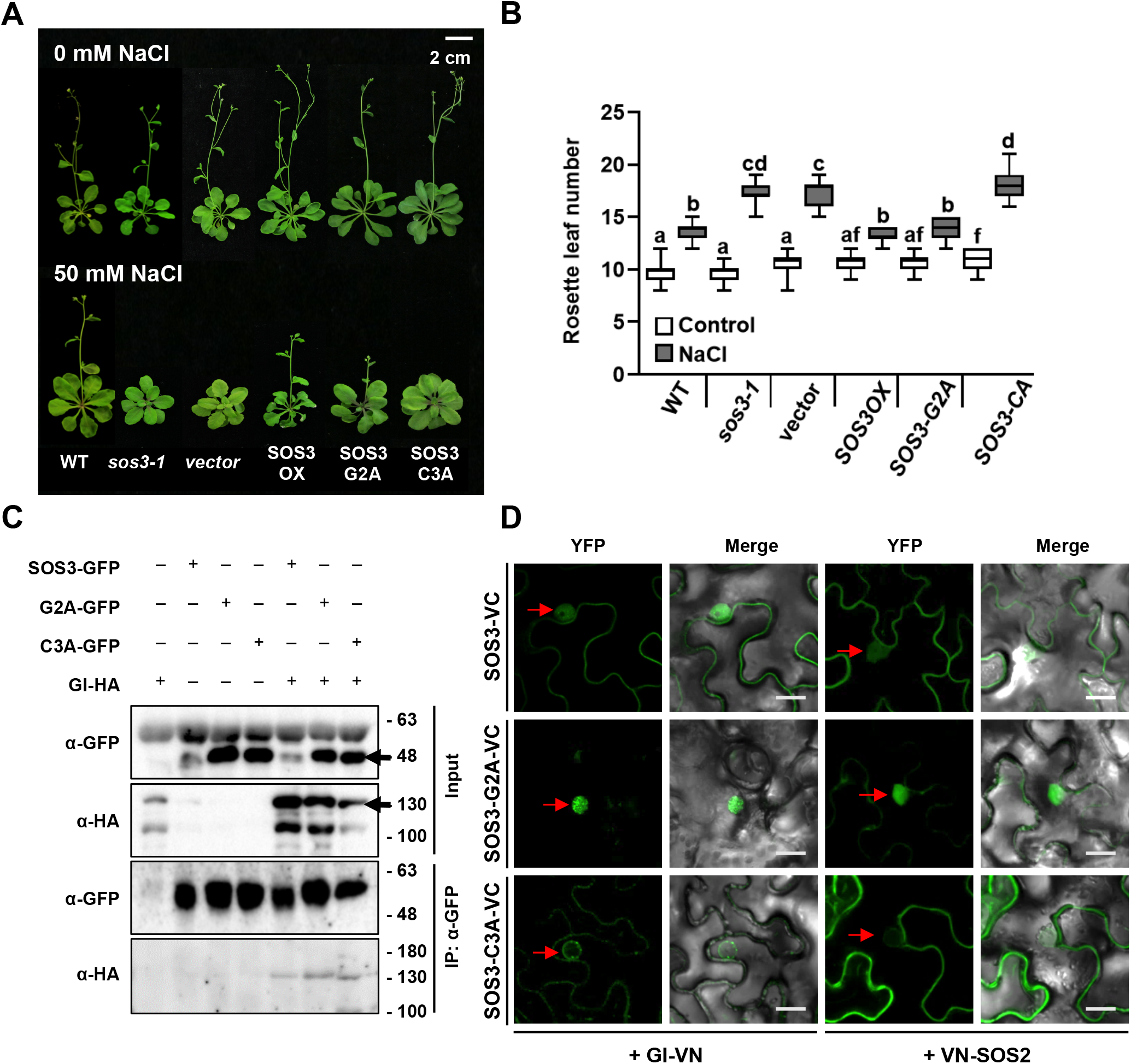
Nuclear localization of SOS3 is required for flowering under salt stress. (**A**) Effect of salt on the flowering time of wild-type (WT), mutant *sos3-1*, and the *sos3-1* mutant expressing the wild-type (SOS3-OX), or non-acylated proteins SOS3-G2A and SOS3-C3A. Eight-day old seedlings were transferred to MS media supplemented with 50 mM NaCl. Photographs were taken after bolting. (**B**) Rosette leaf number was counted at bolting as flowering time. Shown are the Box plots: center lines show the medians; box limits indicate the 25th and 75th percentiles; whiskers extend to the minimum and maximum (n=15). Letters indicate significantly different means at p<0.01, by Fisher’s LSD test. (**C**) Co-immunoprecipitation of GI and SOS3 proteins. Tagged proteins SOS3-GFP, SOS3-G2A-GFP and SOS3-C3A-GFP were transiently co-expressed with GI-HA in tobacco leaves. Total proteins were extracted and immunoprecipitation was done with GFP antibodies (α-GFP). Arrows indicate the target proteins. (**D**) BiFC of mutant SOS3 proteins with GI (left), or SOS2 (right). Indicated proteins were transiently expressed in tobacco leaves. YFP signals were detected under confocal microscope. Scale Bar represents 20 µm.

**Figure 6.**
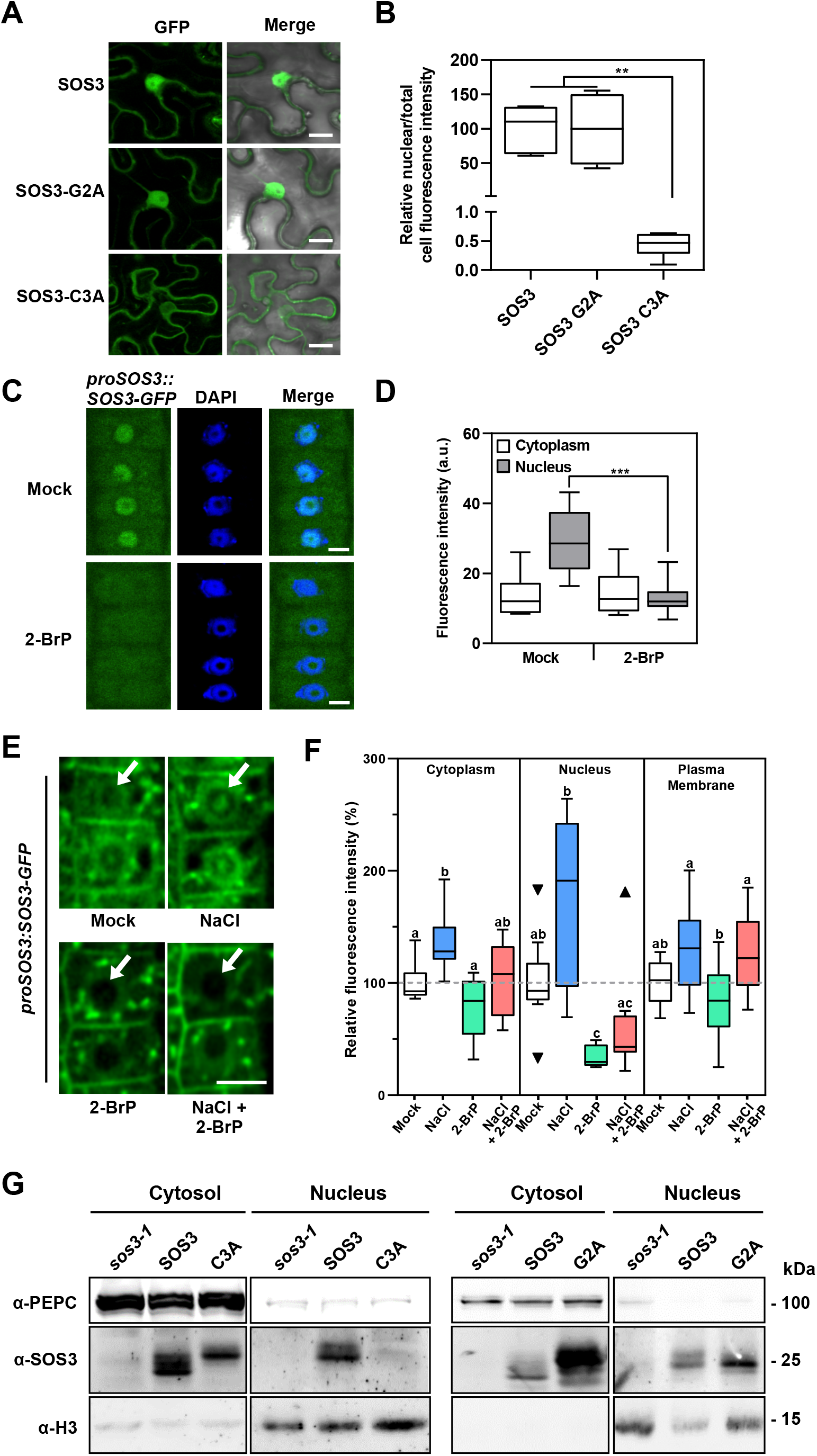
Palmitoylation directs SOS3 nuclear import. (**A**, **B**) Mutation of the *S*-acylation site in SOS3 abrogates import into the nucleus. (**A**) Transient expression in *Nicotiana* of GFP-fused to the C-terminal part of SOS3, SOS3-G2A, and SOS3-C3A was detected under a confocal microscope. Scale bar represents 20 µm. (**B**) Normalized nuclear fluorescence intensity vs. total cell fluorescence. Shown in the Box plot: center lines show the medians; box limits indicate the 25th and 75th percentiles; whiskers extend to the minimum and maximum (n≥3). Asterisks indicate significantly different means of samples with and without salt for each genotype at p<0.01 by Fisher’s LSD test. (**C**, **D**) 2-BrP inhibits SOS3 import to the nucleus. (**C**) Representative images of root meristematic epidermal cells of Arabidopsis *sos3-1* seedlings expressing *proSOS3:SOS3-GFP.* After 5 days growing under control conditions, plants were exposed for one additional day to mock or 50 µM of 2-BrP. Plants were treated with 0.1 % Triton X-100 before imaging to allow counter-staining of nuclei with DAPI. Scale bar 5 µm. (**D**) SOS3-GFP fluorescence signal (mean gray intensity) in cytoplasm and nucleus of cells as shown in (C). Data are represented as in (B) from 13 different plants, 10 cells each. Asterisks indicates means statistically different at *p*<0,001 Fisher’s LSD test. (**E**) Representative fluorescence images of root meristematic epidermal cells of *sos3-1* seedlings expressing *proSOS3:SOS3-GFP* under spinning disc confocal microscopy. Five days old seedlings were exposed for one additional day to the indicated treatments (mock, 100 mM NaCl, 50 µM 2-BrP, and 100 mM NaCl plus 50 µM 2-BrP). Arrows indicate the nuclei. Scale bar is 10 µm. (**F**) Percentage of fluorescence intensity (mean gray intensity) after treatments normalized to the signal in the same cellular compartment under control conditions (mock) of samples shown in (E); Shown are the Box plots based on Tukey methods. Data are from 9 different plants, 10 cells each; Letters indicate significantly different means, based on One-way analysis of variance followed by Tukey’s multiple comparisons test, Dunnett’s T3 test when variance was unequal (nucleus), *p*<0.05. Triangles represent outlier data points, not excluded for statistical analyses. (**G**) Nucleo-cytoplasmic fractionation. Nuclear and cytoplasmic proteins of *sos3-1* plants overexpressing wild-type SOS3, or mutants G2A (right panel) and C3A (left), were fractionated and probed with antibodies against SOS3, the cytoplasmic marker protein PEPC, and the nuclear marker Histone3 (H3). Loading of nuclear proteins was 10-fold higher than of cytosolic proteins to compensate for the lower abundance of SOS3 in the nucleus.

To further confirm the S-acylation-dependent nuclear import of SOS3 in Arabidopsis, the *sos3-1* mutant was transformed with the construct *proSOS3:SOS3-GFP*, comprising a genomic copy of the *SOS3* gene to which GFP was added in frame, and designed to mimic the native *SOS3* gene expression. Treatment of these transgenics with the potent palmitoyl-transferase inhibitor 2-bromo-palmitate (2-BrP) resulted in the complete exclusion of SOS3 from the nucleus (Figure 6C-D). The nuclear integrity was not noticeably affected by 2-BrP, as revealed by DAPI staining. Counter-staining with DAPI to visualize the nucleus under regular confocal microscopy required the co-treatment with Triton-X100 to permeate the dye, but the detergent removed the SOS3-GFP signal at the plasma membrane. Therefore, we used spinning-disc confocal laser microscopy (SDCLM) to measure the relative amounts of SOS3-GFP at the plasma membrane, cytoplasm (comprising cytosol and endosomes) and nuclei (Figure 6E-F). In SDCLM, multiplex laser excitation allows detection of the emission light at multiple points simultaneously for high-speed image acquisition and enhanced sensitivity towards low-abundance fluorescent proteins. Treatment with 2-BrP produced a statistically significant reduction in the nuclear pool of SOS3. Salinity (100 mM NaCl, 1 d) increased the abundance of SOS3 in all compartments, but proportionally more in nuclei (Figure 6E-F). The inhibitory effect of 2-BrP on the nuclear localization of SOS3 dominated over the stimulation by the saline treatment. Last, the nucleo-cytoplasmic partition of SOS3 was inspected in *sos3-1* plants transformed to express the SOS3 protein with and without mutations G2A and C3A. Western blots with SOS3 antibodies of fractionated nuclear and cytoplasmic protein extracts demonstrated that protein SOS3-C3A was excluded from nucleus whereas the non-myristoylated SOS3-G2A mutant protein (which was still S-acylated; see Figure 4) was imported into the nucleus (Figure 6G). Together, these data are evidence of the salinity-induced and S-acylation-dependent nuclear import of SOS3.

Next, we tested whether the S-acylation and nuclear import of SOS3 also had a function in salt tolerance. Contrary to the wild-type SOS3, the SOS3-G2A mutant failed to suppress the salt sensitivity of *sos3-1* plants (Supplemental Figure S6), confirming that myristoylation is essential for SOS3 function in salinity tolerance (Ishitani et al., 2000; Quintero et al., 2002). Notably, SOS3-C3A largely rescued the hypersensitivity of *sos3-1*, implying that S-acylation of SOS3 is not required for salt tolerance and is only critical for flowering under salt stress.

### SOS3 interacts with GI and FKF1 to regulate *CO* expression under salt stress

FKF1, ZTL/LKP1 and LKP2 are blue-light photoreceptors that mediate light-dependent protein degradation of floral regulators by the E3 ubiquitin ligase complex (Zoltowski and Imaizumi, 2014). FKF1 associates with GI to degrade CDF1, a *CO* transcriptional repressor that acts in late afternoon in LDs (Imaizumi et al., 2005; Sawa et al., 2007; Suarez-Lopez et al., 2001). Salt induced degradation affected GI but not FKF1 since the abundance of neither the *FKF1* transcript nor the FKF1 protein changed significantly under salt treatment (Supplemental Figures S1 and S7). To test whether nuclear-imported SOS3 associates with the GI-FKF1 complex, tagged proteins GI-HA, FKF1-MYC and SOS3-FLAG were co-expressed in tobacco leaves and submitted to co-IP with anti-FLAG antibodies. SOS3 pulled down both GI and FKF1 under regular and saline conditions (Figure 7A), suggesting that SOS3 does interact with the GI-FKF1 complex. Salt did not affect the interaction of GI and FKF1 (Figure 7A).

**Figure 7.**
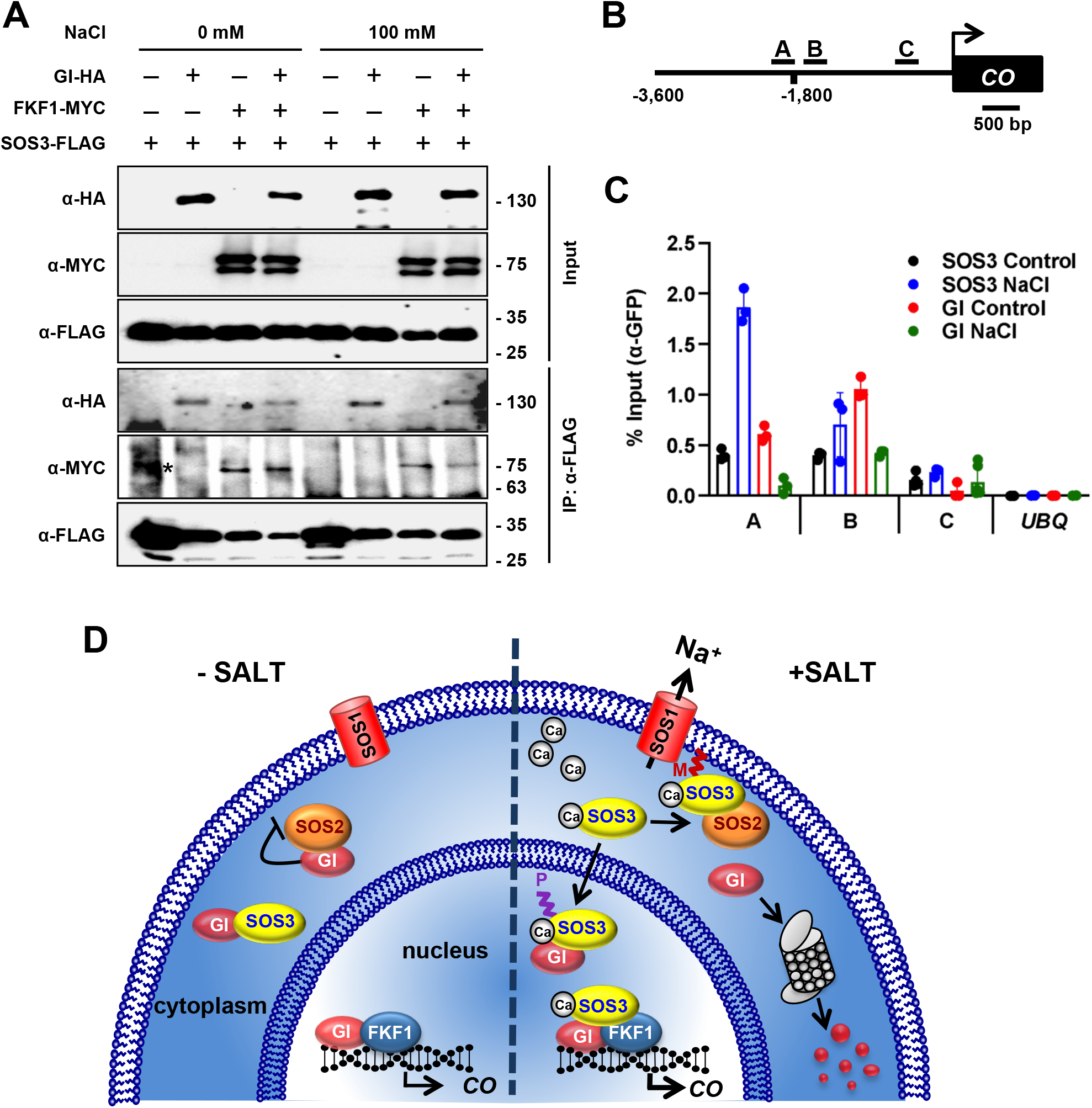
SOS3 forms a complex with GI and FKF at the *CO* promoter. (**A**) SOS3-FLAG was transiently co-expressed with GI-HA and/or FKF1-MYC in tobacco leaves. Total proteins from leaves treated with 100 mM NaCl for 8 h were extracted and immunoprecipitated with FLAG antibodies (α-FLAG). *, non-specific band. (**B**,**C**) Schematic drawing of the *CO* gene promoter, and locations of amplicons (A, B, and C) for ChIP analysis. (**C**) Salt-induced association of SOS3 onto *CO* promoter. Chromatin isolated from two-week old *GI-GFPox* (GI) and *SOS3-GFPox* (SOS3) plants treated (NaCl) with 100 mM NaCl for 10 h or not (Control), was immunoprecipitated with α-GFP antibodies. Immunoprecipitated and input DNA were used as templates for qPCR using primers specifically targeting to the amplicons A, B, and C. *UBQ10* was used as control. Data is fragment enrichment as percent of input DNA. Error bars represent SE (n≥2). The experiment was repeated two times with similar results. (**D**) Simplified working model: Upon salt stress, SOS3 senses and binds elevated cytosolic Ca^2+^. Calcium-bound SOS3 activates and recruits SOS2 to the plasma membrane through the myristoylation of SOS3, to phosphorylate and activate SOS1, a Na^+^ transporter mediating Na^+^ exclusion and salt tolerance. Free cytosolic GI is degraded to delay flowering. S-acylated SOS3 enters the nucleus to make a transcriptional complex with GI and FKF1 that supports *CO* expression to ensure later flowering under salt stress.

ZTL/LKP1 and LKP2 appear to have functions different to FKF1 in photoperiodic flowering. Similar to *gi* mutants, the *fkf1* mutant flowers late (Nelson et al., 2000), whereas *ztl* and *lkp2* mutants show a wild-type flowering pattern (Imaizumi et al., 2005; Somers et al., 2004). However, plants that overexpress *ZTL* and *LKP2* exhibit late flowering under LDs due to the low expression of *CO* and *FT* (Kiyosue and Wada, 2000; Somers et al., 2004). Whereas salt-induced flowering delay was suppressed in *fkf1*, which had unconditional late flowering, mutants *lkp2* and *ztl103* flowered at a similar time than wild-type plants under both normal and salt stress conditions (Supplemental Figure S8). Further, the phenotype of the double mutants *fkf1 ztl103* and *ztl103 lkp2*, and the triple mutant *fkf1 lkp2 ztl103* indicated that only FKF1 among the blue-light receptors regulates the salt-induced delay in flowering. Last, when wild-type, *gi-2*, *fkf1*, *lkp2*, and *ztl103* plants were treated with salt, only mutations of *gi-2* and *fkf1* conferred salt tolerance (Supplemental Figure S9). Together these data indicate that FKF1, specifically among other E3 ligase photoreceptors, acts at the interface between salt stress response and time of flowering signaling.

The FKF1-GI complex associates with the *CO* promoter to induce flowering (Sawa et al., 2007), and our evidence that SOS3 co-IPed with these proteins suggested that SOS3 could be present at the transcriptional complex regulating *CO* transcription. A chromatin-immunoprecipitation (ChIP) analysis of *sos3-1* transgenic plants expressing *SOS3-GFP* showed the enrichment of SOS3-GFP in amplicons A and B where GI and FKF1 associate most with the *CO* promoter (Figure 7B and Supplemental Figure S10) (Sawa et al., 2007). Fragment C of the *CO* promoter not binding GI (Sawa et al., 2007) and the *UBQ10* promoter were used as negative controls (Figure 7B-C and Supplemental Figure S10). SOS3-GFP association with the *CO* promoter increased 4 to 5-fold upon NaCl treatment (Figure 7C and Supplemental Figure S10). By contrast, the saline treatment reduced GI abundance in the *CO* promoter, reflecting the instability of GI under these stress conditions (Figure 7C). Together, these results indicate that S-acylation enables nuclear translocation of SOS3 to associate with the GI-FKF complex to enhance *CO* expression and promote flowering upon salt stress.

## DISCUSSION

### Salt stress and flowering time

Plants adjust their transition from vegetative growth to reproduction by constantly monitoring and integrating environmental cues. Water or nutrient deprivation often leads to earlier flowering presumably because the lack of essential resources inevitably halts growth, whereas a transitory or mild stress is likely to postpone flowering so that reproduction can resume at a later time (Kazan and Lyons, 2016; Maggio et al., 2018). Stress-induced early flowering is an emergency response to proceed to the next generation when vegetative plants cannot cope with adverse environmental conditions (Takeno, 2016). For instance, the drought-escape response entails adaptive shortening of the vegetative growth phase and anticipated seed production before severe dehydration becomes lethal (Riboni et al., 2013). By contrast, salinity delays flowering in several species, including Arabidopsis (Kim et al., 2007; Kim et al., 2013a; Li et al., 2007; Ryu et al., 2014). Plausibly, this reproductive strategy reflects that non-lethal saline levels reduce but do not impede vegetative growth, since plants have developed adaptive strategies to overcome both the osmotic and ionic stresses imposed by salinity (Munns and Tester, 2008). In this regard, the ubiquitous SOS pathway enables plants to deal with excess Na^+^ through the coordination of ion fluxes back to the soil solution and into the xylem to protect roots from damage (Ji et al., 2013; El Mahi et al., 2019). We suggest that the ecophysiological meaning of salt-induced flowering delay is to allow plants to adapt by simultaneously reducing growth rate and altering the developmental program to extend the vegetative growth phase long enough to gather sufficient metabolic resources to ensure robust flowering and seed filling (Achard et al., 2006; Achard et al., 2007; Wang et al., 2018). From this evolutionary perspective, it is beneficial that the control of flowering time and the physiological response to salinity stress are molecularly linked (Kazan and Lyons, 2016), in this case through the physical interaction and mutual regulation of GI, SOS2 and SOS3, to coordinately mount salt tolerance and postpone reproduction. That gibberellin GA4 counteracted salinity-induced late flowering (Li et al., 2007) supports the notion that delayed flowering is a pro-active, genetically ingrained stress response partly dependent on DELLA proteins (Achard et al., 2006).

### Photoperiod-dependent flowering under saline stress requires GI stabilization by SOS3

Previous studies have shown that salinity-induced delay in the flowering time of Arabidopsis occurs in a dosage dependent manner by reducing transcription of the floral integrators *CO* and *FT* (Kim et al., 2007; Li et al., 2007). The GI protein, a major regulator of photoperiodic-induced flowering through the CO-FT module, also plays a substantial role as a negative regulator in the SOS-mediated salt stress adaptation pathway by sequestering the SOS2 kinase in the cytoplasm (Kim et al., 2013a). Salt induced degradation of GI results in the release of the SOS2 kinase, which in turn makes a complex with SOS3 that is recruited to the plasma membrane for the activation of Na^+^/H^+^ antiporter SOS1 (Kim et al., 2013a; Quintero et al., 2002). The *gi-1* mutant exhibits a de-repressed SOS pathway and exceptional salt tolerance compared to the wild type. Despite the multiplicity of functions of SOS2 in various processes pertinent to adaptation to salinity (Qiu et al., 2004; Cheng et al., 2004; Batelli et al., 2007), SOS2 does not seem to play a significant role in setting the flowering time of Arabidopsis on its own (Li et al., 2007; Kim et al., 2013a). However, our study reveals that SOS3, a critical regulator of SOS2, does modulate the initiation of flowering under salt stress by binding to and stabilizing GI. We show that the salt-dependent GI degradation previously reported mostly occurs in the cytosol, whereas the nuclear pool of GI is preserved by a mechanism that involves its physical interaction with SOS3 (Figure 2 and Supplemental Figure S2). The abundance of the nuclear pool of the GI protein is drastically reduced in salt-treated *sos3-1* plants, which produces a mutated SOS3 protein unable to interact with GI (Figures 2 and 3). Nuclear localization of the GI-SOS3 complex was abolished in plants expressing the non-S-acylatable SOS3-C3A protein that remained outside the nucleus (Figures 5, 6 and Supplemental Figure S5). Only the nucleus-localized GI is competent to promote flowering (Kim et al., 2013b), and thus the salt-induced flowering delay of the *gi-2* mutant was rescued by expression of the *GI-NLS* protein that preferentially partitions into the nucleus (Figure 2). Together, these results indicate that SOS3 promotes the stabilization of nuclear GI during salt stress and explain why *GI* overexpression was unable to rescue the salt-dependent late flowering of *sos3-1* plants (Figure 1).

### S-acylation promotes nuclear import of SOS3

SOS2/CIPK24 and SOS3/CBL4 belong to a large array of Ca^2+^-dependent protein kinase modules comprising CIPK and CBL subunits that associate with variable specificity. Post-translational modifications of the CBL subunits determine the subcellular localization of the CIPK-CBL modules (Batistič et al., 2010; Luan et al., 2002). Dual fatty acid modifications consisting of N-myristoylation and S-acylation that contribute to differential sorting of CBLs are only found in CBL1, SOS3/CBL4, CBL5 and CBL9 among the ten CBL proteins of Arabidopsis (Batistič et al., 2010; Batistič et al., 2008). In dual fatty acid modifications, N-terminal myristoylation often provides anchorage to cell membranes. Indeed, myristoylation of SOS3 at Gly-2 allows the SOS2-SOS3 complex to associate with the plasma membrane and phosphorylate SOS1 to promote Na^+^/H^+^ antiport activity and Na^+^ efflux (Quintero et al., 2002). S-acylation of the cysteine residue adjacent to the myristoylated glycine is thought to enhance the membrane attachment of CBL proteins and to regulate subcellular trafficking from the ER to the plasma membrane (Held et al., 2011; Batistič et al., 2008; Batistič and Kudla, 2004; Saito et al., 2018). For instance, the CBL4-CIPK6 complex modulates the K^+^ channel function of AKT2 by promoting its sorting from the ER to the PM (Held et al., 2011). Mutants in each of the three components of this functional module, *cbl4*, *cipk6* and *akt2*, exhibit delayed flowering only in short-day conditions but not in LDs (Held et al., 2011). The floral regulators that were altered in these mutants were not investigated and thus the precise molecular and biochemical connection between K^+^ status and flowering time remains unknown. Here we show that S-acylation of SOS3 is a requisite for nuclear import to secure flowering under LD and saline conditions. Mutation of the S-acylation site (SOS3-C3A) or biochemical inhibition with 2-BrP impeded translocation to the nucleus (Figure 6), and restricted the interaction of SOS3-C3A with SOS2 and GI to the cytoplasm or the perinuclear rim, respectively (Figure 5D and Supplemental Figure S5). The SOS3-C3A mutant was unable to complement the salt-dependent late flowering of the *sos3-1* mutant, whereas SOS3 and SOS3-G2A localized to the nucleus, interacted with GI and supported flowering. Reciprocally, SOS3-C3A was able to suppress most of the salt-sensitivity of *sos3-1*, whereas the non-myristoylatable mutant SOS3-G2A could not (Supplemental Figure S6). This indicates that S-acylation of SOS3 is specifically important for flowering under salt stress but dispensable for salt tolerance. In conclusion, S-acylation targets SOS3 to the nucleus where it forms a complex with GI and helps initiate flowering, whereas cytoplasmic SOS3 functions in the SOS pathway to help establish salt tolerance (Figure 7D).

Eukaryotic palmitoyl-acyl transferases (PATs) of the DHHC (Asp-His-His-Cys) family catalyze protein S-acylation (Batistič, 2012; Hemsley, 2020). In turn, thioesterases break down the ester bond of S-acylation and release the fatty acid. The unique reversibility of protein S-acylation allows proteins to rapidly change their location between intracellular compartments (Aicart-Ramos et al., 2011; Hemsley, 2020). Conditional S-acylation is known to serve as lipid anchor at membranes to immobilize and restrain proteins from entering the nucleus (Hemsley, 2020; Eisenhaber et al., 2011; Lott et al., 2011). For instance, the osmotic-stress responsive transcription factor NFAT5a is myristoylated, S-acylated and sorted to the plasma membrane of animal cells. Upon osmotic stress, NFAT5a moves into the nucleus by a mechanism likely involving de-S-acylation (Eisenhaber et al., 2011), which is the reverse to the novel mechanism we report here for SOS3 nuclear import. To date there is no molecular mechanism known to control nuclear import of a lipid-modified protein in a regulated manner (Aicart-Ramos et al., 2011; Chamberlain and Shipston, 2015). SOS3 lacks canonical nuclear localization signals, which suggests that SOS3 enters the nucleus assisted by a shuttle or gateway protein, whose interaction is presumably dependent on the S-acylation status of SOS3. The finding that non-S-acylated SOS3 was still able to interact with GI and that the complex was detected at the nuclear rim discards the trivial possibility that GI shuttles SOS3 to the nucleus as a complex. While SOS3 S-acylation is a strict requirement for nuclear recruitment, it remains unknown whether SOS3 undergoes de-S-acylation when entering the nucleus or is processed therein. We posit a model (Figure 7D) in which a fraction of the SOS3 protein is S-acylated and partitioned into the nucleus. Upon salinity stress, a Ca^2+^ spike would activate SOS3, fostering the interaction with GI since Ca^2+^ supplementation strengthened SOS3-GI interaction. Simultaneously, the N-myristoylated but non-S-acylated SOS3 remaining in the cytosol recruits SOS2 to activate SOS1 (Quintero et al., 2002). Quantitative data in Figure 6F indicates that salinity stress enhances S-acylation and transfer of SOS3 to the nuclear pool relative to the whole cell content, a process that was blocked *in vivo* by the PAT inhibitor 2-BrP.

Often, protein S-acylation is concurrent with myristoylation or prenylation because substrate proteins for palmitoyl-S-transferases (PATs) must be attached to or in the vicinity of membranes (Rana et al., 2018). However, N-myristoylation is not a biochemical requirement for eukaryotic PATs since S-acylation can occur throughout the target protein. Indeed, the non-myristoylatable SOS3 mutant G2A, but not the double mutant G2A/C3A, was still S-acylated, in tobacco and yeast (Figure 4). Moreover, the SOS3-G2A protein was readily detected in Arabidopsis nuclei (Figure 6). Understanding the environmental and biochemical inputs that elicit S-acylation (and de-S-acylation) of SOS3, and identify the PATs involved in this process (24 putative *PAT* genes in Arabidopsis) will be a promising line of research. PAT10 functions in salt tolerance, polar growth of root hairs, and stomatal movements in Arabidopsis (Song et al., 2018; Zhou et al., 2013; Zhang et al., 2015), but PAT10 is tonoplast-localized and unlikely to mediate SOS3 palmitoylation. Moreover, palmitoylation of CBL2 and CBL3 by PAT10 results in the attachment of these target CBLs to the tonoplast (Song et al., 2018). The presence of a PAT localized in the nuclear envelope or in the perinuclear ER membrane (Batistič, 2012) and driving the palmitoylation-dependent nuclear import of SOS3 is an attractive possibility.

### SOS3 ensures GI-mediated flowering under salt stress

Salinity stress delays flowering due to GI degradation (Kim et al., 2013a) and the reduced transcription of the floral integrators *CO* and *FT* (Kim et al., 2007; Li et al., 2007). Here we show that nuclear GI is more recalcitrant to degradation than cytosolic GI since the GI-SOS3 complex is stable inside the nucleus. Removal of SOS3 destabilizes nuclear GI and delays flowering even further. Thus, a novel function of SOS3 is to ensure that flowering will occur in a saline environment, albeit at a later time compared to non-stressing conditions. Ultimately, the developmental transition to flowering requires the de-repression of *CO* and *FT*, and GI induces flowering mainly through the CO-FT module (Sawa et al., 2007; Sawa and Kay, 2011). Expression of *CO* under long days requires the degradation of repressors collectively known as CDFs that delay flowering by repressing *CO* transcription. A protein complex formed by GI and FKF1 promotes degradation CDFs (Imaizumi et al., 2005; Sawa et al., 2007; Fornara et al., 2009; Nelson et al., 2000). Similarly to *gi* mutant, plants bearing the *fkf1* mutation lost the salt-induced flowering delay and flowered late unconditionally, with no statistical difference with and without salt stress (Supplemental Figure S8). These mutants also display increased salt tolerance, similar to *gi* mutants (Supplemental Figure S9). Even though SOS3 was able to interact with GI and FKF1 both in normal and saline conditions (Figure 7A), the association of SOS3 to the GI and FKF1 binding sites in the *CO* promoter region increased upon salt treatment, thereby leading to enhanced *CO* expression upon the onset of salt stress (Figure 1C). This result and the spinning disc confocal microscopy data (Figure 6E-F) support the notion that salt stress enhances nuclear import of SOS3 through S-acylation to promote the expression of *CO* (Figures 1 and 7). We have not yet investigated whether CDF repressors are displaced or degraded upon binding of the GI-FKF1-SOS3 complex.

*CO* and *FT* transcripts in *sos3-1* followed wild-type dynamics in non-saline conditions, but salt treatment reduced *CO* and abated *FT* transcript levels in the mutant (Figure 1C), indicating that SOS3 functions to sustain the expression of these two critical floral integrators. Our data confirms previous reports of reduced *FT* expression under salinity but are partially in conflict with the concomitant repression of *CO* that has been reported in wild-type Arabidopsis after several days (5-10 d) under saline stress (Li et al., 2007). However, Kilian et al. (2007) showed that with salt treatment given at ZT3, *CO* expression reached a maximum at ZT15 (dusk), but expression during the dark period was not recorded. Here, we analyzed the diurnal pattern of *CO* and *FT* upon the onset of saline treatment, coinciding with the beginning of the enhanced nuclear import of SOS3 (Figures 6 and 7). *CO* expression and CO protein abundance are known to be under multiple and complex layers of regulation (Shim et al., 2017), and the expression pattern could change along the process of salt-adaptation. Indeed, quantitation of *CO* and *FT* transcripts at 1- and 5-day showed that the salt- and genotype-dependent reduction in *FT* expression was more intense after 5 days in salt compared to 1 day, and that *CO* transcript abundance showed a similar trend after the 5-day treatment (Supplemental Figure S11). The negative impact of the *sos3-1* mutation was observed regardless of the duration of the salt treatment. The early increase in the abundance of *CO* transcript in the wild-type upon salt treatment and the opposing decrease in the *sos3-1* mutant is coherent with the model depicted in Figure 7D. In the wild-type, the nuclear pool of GI is not degraded but protected by the nuclear-imported SOS3 and, together with GI and FKF1, up-regulates *CO* transcription. However, in the *sos3-1* mutant, nuclear GI is also degraded and *CO* transcription is compromised. What ultimately determines floral commitment is FT, and our *FT* expression data is in agreement with delayed flowering in the wild-type and the acute delay in the *sos3-1* mutant (Figure 1). Although CO is a major transcriptional activator of *FT*, GI also promotes *FT* expression in a CO-independent manner (Jung et al., 2007; Sawa and Kay, 2011). Because GI protein abundance decreased the most in the *sos3-1* mutant under salt stress, a condition that reduced *CO* transcription and completely abated *FT* transcripts (Figure 1 and Supplemental Figure S11), it appears that *FT* transcription under saline stress is largely determined by GI abundance and only partly dependent on CO. It remains to be investigated whether the GI-FKF1-SOS3 complex also regulates *FT* transcription directly as with *CO*.

Altogether we have shown that the calcium-binding protein SOS3, that is an upstream regulator of the salt tolerance determinants SOS2 and SOS1, also functions to ensure the completion of flowering under salt stress by stabilizing the nuclear pool of GI and promoting the expression of *CO* (Figure 7D). These results add new layers of regulation and molecular connections to the mechanism that links salt stress adaptation and the photoperiodic flowering pathway. Our results also expand the repertoire of cellular processes in which SOS3 serves as an integrator transmitting environmental inputs leading to stress adaptation and transition to reproductive stage, and uncover a potentially novel mechanism for palmitoylation-dependent ingress of proteins into the nucleus.

## METHODS

### Plant materials and flowering under salinity stress

Mutants *sos1-1, sos2-2, sos3-1, cbl10/scabp8* (SALK_056042), *gi-2*, and transgenic lines overexpressing tagged *GI-HA* (called *GI-OX*) have been described previously (Fowler et al., 1999; Ishitani et al., 2000; Quan et al., 2007). *GIpro:GI-GFP* (called *GI-GFP*), *GIpro:GI-GFP-NLS* (called *GI-NLS*), and *GIpro:GI-GFP-NES* (called *GI-NES*) transgenic plants in *gi-2* mutant background were kindly provided by Hong Gil Nam (Daegu Gyeongbuk Institute of Science and Technology, Korea) (Kim et al., 2013b). Transgenic lines expressing *SOS3*, *SOS3-G2A*, *SOS3-C3A*, and *SOS3-GFP* from the *35S* promoter, and the genomic construct *proSOS3:SOS3-GFP* were generated in the *sos3-1* background by floral dipping (further details are in Supplemental Information, under Plasmid Construction). Mutant lines *fkf1/ztl103, ztl103/lkp2 and fkf1/lkp2/ztl103* were gently provided by Takato Imaizumi (University of Washington, USA) (Baudry et al., 2010). Lines *sos1-1 GI-OX*, *sos2-2 GI-OX*, *sos3-1 GI-OX* and *cbl10 GI-OX* were generated by crossing *GI-OX* transgenics with mutants *sos1-1, sos2-2, sos3-1* and *cbl10.* Plants were confirmed for late flowering and salt sensitivity, and further verified by PCR or western blot for GI expression.

All seeds were sterilized with 70% ethanol and 2% bleach (sodium hypochlorite solution) and stratified at 4°C for 2-3 days. Plants were grown under long-day (LD) conditions (16 h light / 8 h dark, 80-100 µM m^-2^s^-1^) at 23°C. For flowering phenotype seeds were first grown on full MS media (Duchefa, Haarlem, Netherlands) containing 1% sucrose (supplemented with vitamins, 2.5% phytagel) (Murashige and Skoog, 1962). Then 8-day old seedlings were transferred to MS media (basal salt MS without vitamins) with or without NaCl in plant culture dishes (14 cm in height; 10 cm in diameter) (SPL Life Science, Pocheon, Korea). Six plants were planted per plant dish with sufficient air exchange. Salt-treatments were adjusted depending on the genotype and the relative sensitivity or tolerance to salinity of plants used in each experiment, to ensure that plants survived the treatment and flowered. The concentrations of NaCl used are indicated in the Figure legends. Total rosette leaf numbers were counted after bolting to indicate flowering time.

### Bimolecular fluorescence complementation (BIFC)

Plasmid constructs for BiFC were transformed into *Agrobacterium tumefaciens* strain GV3101. Two days after *Agrobacterium* infiltration into tobacco leaves (see Supplemental Methods for details), solutions of 100 mM NaCl or 3 mM CaCl_2_, with or without 2 mM EGTA were infiltrated into tobacco epidermal cells, and 6-8 h later YFP signals were detected under confocal laser scanning microscope (FV 1000 Olympus, Tokyo, Japan). Excitation and emission wavelengths for YFP are 515 nm and 527 nm, respectively. The same settings were used for fluorescence detection in all the samples within the same experiment.

### Immunoblotting and immunoprecipitation

Fused proteins GI-HA, SOS2-GFP, SOS3-MYC and MYC-SOS3-1 were transiently expressed in *N. benthamiana* leaves alone or in given combinations by *Agrobacterium* infiltration. Leaves were treated or untreated with 100 mM NaCl or 3 mM CaCl_2_. EGTA (2 mM) was used as a calcium chelator to inhibit calcium signaling. Immunoblotting followed standard procedures (Kim et al., 2013a). Buffer composition is given in Supplemental Methods.

For immunoprecipitation, rat α-HA (1:250, Roche, #11867423001, Indianapolis, IN) or Rabbit α-GFP polyclonal (1:250, Invitrogen, #A11120) antibodies were pre-incubated with protein A agarose (Invitrogen) at 4°C for 30 min. Protein extracts were added and further incubated for 1 h at 4°C. Complexes were separated by SDS–PAGE. Each immunoblot was incubated with the appropriate primary antibody (α-HA (1:2000), α-GFP (1:5,000, Abcam, #ab6556, Cambridge, MA) or α-MYC (1:1,000, Cell Signaling Technology, #2276, Danvers, MA) for 1 to 2 h at room temperature or overnight at 4°C. The antigen protein was detected by chemiluminescence using an ECL-detecting reagent (Bio-Rad, Hercules, CA) and signals were detected by Imaging system (ChemiDoc^TM^MP, Bio-Rad).

### Nucleus preparation

To test salt-induced degradation of cytosolic and nuclear GI protein, 12-day old Arabidopsis *GI-OX*, *sos3-1 GI-OX* and *sos2-2 GI-OX* were treated with or without 100 mM NaCl in distilled water for 12 h. To determine the subcellular localization of SOS3, SOS3-G2A and SOS3-C3A proteins, aerial parts of 4-week old *sos3-1* plants expressing these proteins were collected for fractionation of nuclear and cytosolic proteins. Nuclei were purified using Plant Nuclei Isolation/extraction Kit (Sigma) and proteins extracted with Laemmli buffer. Cytosolic proteins were obtained by precipitating supernatants of the first nuclei pelleting step with 10% TCA and resuspending in a denaturing buffer consisting of 50 mM Tris-HCl (pH 7.8), 4 M urea, 2% SDS and 2.5% glycerol. Commercially available antibodies against α-Histon3 (Abcam, #ab1791) and α-Phospho Enol Pyruvate Carboxylase (PEPC) (Agrisera, #AS09 458, Vännäs, Sweden) were used as nuclear and cytosolic markers, respectively.

### Detection of SOS3 S-acylation by differential alkylation

Wild-type SOS3 and mutant proteins G2A, C3A and G2A/C3A, all with 6xHis tags, were expressed in yeast. Protein extracts were first treated with N-ethylmaleimide (NEM) to block free cysteine thiols, next with hydroxylamine to break palmitoyl-thioester bonds, and then with methyl-PEG_24_-maleimide, MM(PEG)_24_, to alkylate newly formed cysteine thiols. Chemicals and detailed procedure are in Supplemental Methods. Proteins were resolved in 11% acrylamide SDS-PAGE gels and subjected to western blot analysis using the α-SOS3 antibody (Ishitani et al., 2000) at 1:2000 dilution. The theoretical mass of SOS3 with 6x His tag is 26.5 kDa and MM(PEG)_24_ adds 1.24 kDa for each cysteine alkylated.

### Acyl resin-assisted capture (acyl-RAC) of SOS3

The wild-type SOS3 and mutants G2A, C3A and G2A/C3A were tagged C-terminally with a Tandem Affinity Purification (TAP) tag and expressed in *N. benthamiana* leaves. Proteins extracted from leaf tissue were processed following the acyl-RAC method as described by Chai et al (2019); the detailed procedure is given Supplemental Methods. Before sample processing, aliquots were withdrawn to be analyzed as “input”. Protein samples were divided in two parts and hydroxylamine was added at 0.5 M final concentrations to one of these parts to break *S*-acyl-thioester bonds, whereas the other served as untreated control. Each sample was incubated with thiopropyl-sepharose 6B resin (Sigma) to link proteins with free thiols. Protein eluted with a DTT-containing buffer were analyzed by western blotting with the α-SOS3 antibody.

### 2-Bromo-palmitate treatment and microscopy

Plants were sown in ½ MS plates with 1% sucrose in a LD chamber at 21°C. Five days after germination the plant were incubated in liquid in ½ MS with 1% sucrose media with or without 50 µM 2-bromopalmitate (2-BrP; Sigma-Aldrich), and with or without 100 mM NaCl, for 24 h keeping the same growth conditions. For controls, a mock treatment with the same volume of ethanol (2-BrP solvent) was added to samples without 2-BrP. Details about treatments and microscopy are given in Supplemental Methods.

### RNA isolation and Q-RT PCR

Gene expression was analyzed by quantitative RT-PCR as detailed in Supplemental Methods. Each data point represents the average of three independent amplifications of the same RNA sample run in the same reaction plate. Each biological replicate had three technical replicas. Primers used for Q-RT PCR are in Supplemental Table S1.

### Chromatin immunoprecipitation (ChIP) assay

Two-week old Arabidopsis seedlings (*GI-GFPox* and *SOS3-GFPox*) treated with 100 mM NaCl for 10 h were used for the ChiP assay. Procedures of fixation and isolation of chromatin were performed as described (Sawa et al., 2007; Saleh et al., 2008). Detailed description can be found in Supplemental Methods.

### Statistical analyses

The statistical analysis used to obtain the significance level is indicated in the legend to each figure. The different statistical analyses were performed using GraphPad Prism version 8, GraphPad Software, San Diego, California USA, www.graphpad.com.

## AUTHOR CONTRIBUTIONS

H.J. Park, R. Aman and C.J. Lim performed the experiments related to flowering time and protein interactions. F. Gámez, I. Villalta, E. Garcia, M. Lindahl, R. Carranco, and F.J. Quintero determined the acylation, subcellular localization and the salt-tolerance function of wild-type and mutant SOS3 proteins. H.J. Park, F. Gámez, C. Sánchez-Rodríguez, F.J. Quintero, J.M. Pardo, and W.-Y. Kim designed experiments and analyzed data. R.A. Bressan and S.Y. Lee discussed data. H.J. Park, F. Gámez, J.M. Pardo, F.J. Quintero and D.J. Yun wrote the manuscript. J.M. Pardo, W.-Y. Kim, C. Sánchez-Rodríguez, F.J. Quintero and D.-J. Yun supervised the project.

## ACKNOWLEDGMENT

We thank Hong Gil Nam (Daegu Gyeongbuk Institute of Science and Technology, Korea) and Takato Imaizumi (University of Washington, USA) for *GIpro:GI-GFP* (called *GI-GFP*), *GIpro:GI-GFP-NLS* (called *GI-NLS*), and *GIpro:GI-GFP-NES* (called *GI-NES*) transgenic plant seeds, and *fkf1/ztl103, ztl103/lkp2 and fkf1/lkp2/ztl103* mutant seeds, respectively.

## References

Achard, P., Cheng, H., De Grauwe, L., Decat, J., Schoutteten, H., Moritz, T., Van Der Straeten, D., Peng, J., and Harberd, N. P. (2006). Integration of plant responses to environmentally activated phytohormonal signals. Science 311:91–94.

Achard, P., Baghour, M., Chapple, A., Hedden, P., Van Der Straeten, D., Genschik, P., Moritz, T., and Harberd, N. P. (2007). The plant stress hormone ethylene controls floral transition via DELLA-dependent regulation of floral meristem-identity genes. Proc. Natl. Acad. Sci. 104:6484–6489.

Aicart-Ramos, C., Valero, R. A., and Rodriguez-Crespo, I. (2011). Protein palmitoylation and subcellular trafficking. Biochim. Biophys. Acta BBA -Biomembr. 1808:2981–2994.

Barragán, V., Leidi, E. O., Andrés, Z., Rubio, L., De Luca, A., Fernández, J. A., Cubero, B., and Pardo, J. M. (2012). Ion exchangers NHX1 and NHX2 mediate active potassium uptake into vacuoles to regulate cell turgor and stomatal function in Arabidopsis. Plant Cell 24:1127–1142.

Batelli, G., Verslues, P. E., Agius, F., Qiu, Q., Fujii, H., Pan, S., Schumaker, K. S., Grillo, S., and Zhu, J.-K. (2007). SOS2 promotes salt tolerance in part by interacting with the vacuolar H^+^-ATPase and upregulating its transport activity. Mol. Cell. Biol. 27:7781–7790.

Batistič, O. (2012). Genomics and localization of the Arabidopsis DHHC-Cysteine-Rich Domain S-Acyltransferase protein family. Plant Physiol. 160:1597–1612.

Batistič, O., and Kudla, J. (2004). Integration and channeling of calcium signaling through the CBL calcium sensor/CIPK protein kinase network. Planta 219:915–924.

Batistič, O., Sorek, N., Schültke, S., Yalovsky, S., and Kudla, J. (2008). Dual fatty acyl modification determines the localization and plasma membrane targeting of CBL/CIPK Ca^2+^ signaling complexes in Arabidopsis. Plant Cell 20:1346–1362.

Batistič, O., Waadt, R., Steinhorst, L., Held, K., and Kudla, J. (2010). CBL-mediated targeting of CIPKs facilitates the decoding of calcium signals emanating from distinct cellular stores. Plant J. 61:211–222.

Baudry, A., Ito, S., Song, Y. H., Strait, A. A., Kiba, T., Lu, S., Henriques, R., Pruneda-Paz, J. L., Chua, N.-H., Tobin, E. M., et al. (2010). F-Box Proteins FKF1 and LKP2 Act in Concert with ZEITLUPE to Control Arabidopsis Clock Progression. Plant Cell 22:606–622.

Brunelli, J. P., and Pall, M. L. (1993). A series of yeast shuttle vectors for expression of cDNAs and other DNA sequences. Yeast 9:1299–1308.

Cao, S., Ye, M., and Jiang, S. (2005). Involvement of GIGANTEA gene in the regulation of the cold stress response in Arabidopsis. Plant Cell Rep. 24:683–690.

Cerdan, P. D., and Chory, J. (2003). Regulation of flowering time by light quality. Nature 423:881–885.

Chai, S., Ge, F.-R., Zhang, Y., and Li, S. (2019). S-acylation of CBL10/SCaBP8 by PAT10 is crucial for its tonoplast association and function in salt tolerance. J. Integr. Plant Biol. 62:718–722.

Chamberlain, L. H., and Shipston, M. J. (2015). The physiology of protein S-acylation. Physiol. Rev. 95:341–376.

Cheng, N.-H., Pittman, J. K., Zhu, J.-K., and Hirschi, K. D. (2004). The protein kinase SOS2 activates the Arabidopsis H^+^/Ca^2+^ antiporter CAX1 to integrate calcium transport and salt tolerance. J. Biol. Chem. 279:2922–2926.

Corbesier, L., and Coupland, G. (2006). The quest for florigen: a review of recent progress. J. Exp. Bot. 57:3395–3403.

Dalchau, N., Baek, S. J., Briggs, H. M., Robertson, F. C., Dodd, A. N., Gardner, M. J., Stancombe, M. A., Haydon, M. J., Stan, G.-B., Gonçalves, J. M., et al. (2011). The circadian oscillator gene GIGANTEA mediates a long-term response of the *Arabidopsis thaliana* circadian clock to sucrose. Proc. Natl. Acad. Sci. 108:5104–5109.

David, K. M., Armbruster, U., Tama, N., and Putterill, J. (2006). Arabidopsis GIGANTEA protein is post-transcriptionally regulated by light and dark. FEBS Lett. 580:1193–1197.

Eisenhaber, B., Sammer, M., Lua, W. H., Benetka, W., Liew, L. L., Yu, W., Lee, H. K., Koranda, M., Eisenhaber, F., and Adhikari, S. (2011). Nuclear import of a lipid-modified transcription factor: Mobilization of NFAT5 isoform a by osmotic stress. Cell Cycle 10:3897–3911.

El Mahi, H., Pérez-Hormaeche, J., De Luca, A., Villalta, I., Espartero, J., Gámez-Arjona, F., Fernández, J. L., Bundó, M., Mendoza, I., Mieulet, D., et al. (2019). A critical role of sodium flux via the plasma membrane Na^+^/H^+^ exchanger SOS1 in the salt tolerance of rice. Plant Physiol. 180:1046–1065.

Fornara, F., Panigrahi, K. C. S., Gissot, L., Sauerbrunn, N., Rühl, M., Jarillo, J. A., and Coupland, G. (2009). Arabidopsis DOF transcription factors act redundantly to reduce CONSTANS expression and are essential for a photoperiodic flowering response. Dev. Cell 17:75–86.

Fornara, F., de Montaigu, A., Sánchez-Villarreal, A., Takahashi, Y., Ver Loren van Themaat, E., Huettel, B., Davis, S. J., and Coupland, G. (2015). The GI–CDF module of Arabidopsis affects freezing tolerance and growth as well as flowering. Plant J. 81:695–706.

Fowler, S., Lee, K., Onouchi, H., Samach, A., Richardson, K., Morris, B., Coupland, G., and Putterill, J. (1999). GIGANTEA: a circadian clock-controlled gene that regulates photoperiodic flowering in Arabidopsis and encodes a protein with several possible membrane-spanning domains. EMBO J. 18:4679–4688.

Greenham, K., and McClung, C. R. (2015). Integrating circadian dynamics with physiological processes in plants. Nat. Rev. Genet. 16:598–610.

Han, Y., Zhang, X., Wang, Y., and Ming, F. (2013). The Suppression of WRKY44 by GIGANTEA-miR172 Pathway Is Involved in Drought Response of *Arabidopsis thaliana*. PLOS One 8:e73541.

Held, K., Pascaud, F., Eckert, C., Gajdanowicz, P., Hashimoto, K., Corratgé-Faillie, C., Offenborn, J. N., Lacombe, B., Dreyer, I., Thibaud, J.-B., et al. (2011). Calcium-dependent modulation and plasma membrane targeting of the AKT2 potassium channel by the CBL4/CIPK6 calcium sensor/protein kinase complex. Cell Res. 21:1116–1130.

Hemsley, P. A. (2020). S-acylation in plants: an expanding field. Biochem. Soc. Trans. 48:529–536.

Imaizumi, T., Schultz, T. F., Harmon, F. G., Ho, L. A., and Kay, S. A. (2005). FKF1 F-Box protein mediates cyclic degradation of a repressor of CONSTANS in Arabidopsis. Science 309:293–297.

Ishitani, M., Liu, J., Halfter, U., Kim, C.-S., Shi, W., and Zhu, J.-K. (2000). SOS3 function in plant salt tolerance requires N-myristoylation and calcium binding. Plant Cell 12:1667–1677.

Ji, H., Pardo, J. M., Batelli, G., Van Oosten, M. J., Bressan, R. A., and Li, X. (2013). The Salt Overly Sensitive (SOS) pathway: Established and emerging roles. Mol. Plant 6:275–286.

Jung, J.-H., Seo, Y.-H., Seo, P. J., Reyes, J. L., Yun, J., Chua, N.-H., and Park, C.-M. (2007). The GIGANTEA-regulated microRNA172 mediates photoperiodic flowering independent of CONSTANS in Arabidopsis. Plant Cell 19:2736–2748.

Kazan, K., and Lyons, R. (2016). The link between flowering time and stress tolerance. J. Exp. Bot. 67:47–60.

Kilian, J., Whitehead, D., Horak, J., Wanke, D., Weinl, S., Batistic, O., D’Angelo, C., Bornberg-Bauer, E., Kudla, J., and Harter, K. (2007). The AtGenExpress global stress expression data set: protocols, evaluation and model data analysis of UV-B light, drought and cold stress responses. Plant J. 50:347–363.

Kim, S.-G., Kim, S.-Y., and Park, C.-M. (2007). A membrane-associated NAC transcription factor regulates salt-responsive flowering via FLOWERING LOCUS T in Arabidopsis. Planta 226:647–654.

Kim, W.-Y., Ali, Z., Park, H. J., Park, S. J., Cha, J.-Y., Perez-Hormaeche, J., Quintero, F. J., Shin, G., Kim, M. R., Qiang, Z., et al. (2013a). Release of SOS2 kinase from sequestration with GIGANTEA determines salt tolerance in Arabidopsis. Nat. Commun. 4:1352.

Kim, Y., Han, S., Yeom, M., Kim, H., Lim, J., Cha, J.-Y., Kim, W.-Y., Somers, D. E., Putterill, J., Nam, H. G., et al. (2013b). Balanced nucleocytosolic partitioning defines a spatial network to coordinate circadian physiology in plants. Dev. Cell 26:73–85.

Kiyosue, T., and Wada, M. (2000). LKP1 (LOV kelch protein 1): a factor involved in the regulation of flowering time in Arabidopsis. Plant J. 23:807–815.

Lakatos, L., Szittya, G., Silhavy, D., and Burgyán, J. (2004). Molecular mechanism of RNA silencing suppression mediated by p19 protein of tombusviruses. EMBO J. 23:876–884.

Li, K., Wang, Y., Han, C., Zhang, W., Jia, H., and Li, X. (2007). GA signaling and CO/FT regulatory module mediate salt-induced late flowering in *Arabidopsis thaliana*. Plant Growth Regul. 53:195–206.

Liu, J., and Zhu, J.-K. (1998). A calcium sensor homolog required for plant salt tolerance. Science 280:1943–1945.

Lott, K., Bhardwaj, A., Sims, P. J., and Cingolani, G. (2011). A minimal nuclear localization signal (NLS) in human phospholipid scramblase 4 that binds only the minor NLS-binding site of importin α1. J. Biol. Chem. 286:28160–28169.

Luan, S., Kudla, J., Rodriguez-Concepcion, M., Yalovsky, S., and Gruissem, W. (2002). Calmodulins and Calcineurin B–like proteins: Calcium sensors for specific signal response coupling in plants. Plant Cell 14:s389–s400.

Maggio, A., Bressan, R., Zhao, Y., Park, J., and Yun, D.-J. (2018). It’s hard to avoid avoidance: Uncoupling the evolutionary connection between plant growth, productivity and stress “tolerance.” Int. J. Mol. Sci. 19:3671.

Mathieu, J., Yant, L. J., Mürdter, F., Küttner, F., and Schmid, M. (2009). Repression of flowering by the miR172 target SMZ. PLoS Biol. 7:e1000148.

Munns, R., and Tester, M. (2008). Mechanisms of salinity tolerance. Annu. Rev. Plant Biol. 59:651–681.

Murashige, T., and Skoog, F. (1962). A revised medium for rapid growth and bioassays with tobacco tissue cultures. Physiol. Plant 15:473–497.

Nelson, D. C., Lasswell, J., Rogg, L. E., Cohen, M. A., and Bartel, B. (2000). FKF1, a clock-controlled gene that regulates the transition to flowering in Arabidopsis. Cell 101:331–340.

Oliverio, K. A., Crepy, M., Martin-Tryon, E. L., Milich, R., Harmer, S. L., Putterill, J., Yanovsky, M. J., and Casal, J. J. (2007). GIGANTEA regulates Phytochrome A-mediated photomorphogenesis independently of its role in the circadian clock. Plant Physiol.144:495–502.

Park, H. J., Kim, W.-Y., Pardo, J. M., and Yun, D.-J. (2016). Chapter eight - Molecular interactions between flowering time and abiotic stress pathways. In International Review of Cell and Molecular Biology (ed. Kwang W. Jeon and Lorenzo Galluzzi, pp. 371–412. Academic Press.

Qiu, Q.-S., Guo, Y., Dietrich, M. A., Schumaker, K. S., and Zhu, J.-K. (2002). Regulation of SOS1, a plasma membrane Na^+^/H^+^ exchanger in *Arabidopsis thaliana*, by SOS2 and SOS3. Proc. Natl. Acad. Sci. 99:8436–8441.

Qiu, Q.-S., Guo, Y., Quintero, F. J., Pardo, J. M., Schumaker, K. S., and Zhu, J.-K. (2004). Regulation of vacuolar Na^+^/H^+^ exchange in *Arabidopsis thaliana* by the Salt-Overly-Sensitive (SOS) pathway. J. Biol. Chem. 279:207–215.

Quan, R., Lin, H., Mendoza, I., Zhang, Y., Cao, W., Yang, Y., Shang, M., Chen, S., Pardo, J. M., and Guo, Y. (2007). SCABP8/CBL10, a putative calcium sensor, interacts with the protein kinase SOS2 to protect Arabidopsis shoots from salt stress. Plant Cell 19:1415–1431.

Quintero, F. J., Ohta, M., Shi, H., Zhu, J.-K., and Pardo, J. M. (2002). Reconstitution in yeast of the Arabidopsis SOS signaling pathway for Na^+^ homeostasis. Proc. Natl. Acad. Sci. 99:9061–9066.

Quintero, F. J., Martinez-Atienza, J., Villalta, I., Jiang, X., Kim, W.-Y., Ali, Z., Fujii, H., Mendoza, I., Yun, D.-J., Zhu, J.-K., et al. (2011). Activation of the plasma membrane Na/H antiporter Salt-Overly-Sensitive 1 (SOS1) by phosphorylation of an auto-inhibitory C-terminal domain. Proc. Natl. Acad. Sci. 108:2611–2616.

Rana, M. S., Kumar, P., Lee, C.-J., Verardi, R., Rajashankar, K. R., and Banerjee, A. (2018). Fatty acyl recognition and transfer by an integral membrane S-acyltransferase. Science 359:eaao6326.

Riboni, M., Galbiati, M., Tonelli, C., and Conti, L. (2013). GIGANTEA enables drought escape response via abscisic acid-dependent activation of the florigens and SUPPRESSOR OF OVEREXPRESSION OF CONSTANS1. Plant Physiol. 162:1706–1719.

Ryu, J. Y., Lee, H.-J., Seo, P. J., Jung, J.-H., Ahn, J. H., and Park, C.-M. (2014). The Arabidopsis floral repressor BFT delays flowering by competing with FT for FD binding under high salinity. Mol. Plant 7:377–387.

Saito, S., Hamamoto, S., Moriya, K., Matsuura, A., Sato, Y., Muto, J., Noguchi, H., Yamauchi, S., Tozawa, Y., Ueda, M., et al. (2018). N-myristoylation and S-acylation are common modifications of Ca^2+^-regulated Arabidopsis kinases and are required for activation of the SLAC1 anion channel. New Phytol. 218:1504–1521.

Sawa, M., and Kay, S. A. (2011). GIGANTEA directly activates Flowering Locus T in *Arabidopsis thaliana*. Proc. Natl. Acad. Sci. 108:11698–11703.

Sawa, M., Nusinow, D. A., Kay, S. A., and Imaizumi, T. (2007). FKF1 and GIGANTEA complex formation is required for day-length measurement in Arabidopsis. Science 318:261–265.

Seo, P. J., and Mas, P. (2015). STRESSing the role of the plant circadian clock. Trends Plant Sci. 20:230–237.

Shim, J. S., Kubota, A., and Imaizumi, T. (2017). Circadian clock and photoperiodic flowering in Arabidopsis: CONSTANS is a hub for signal integration. Plant Physiol. 173:5.

Somers, D. E., Kim, W.-Y., and Geng, R. (2004). The F-Box protein ZEITLUPE confers dosage-dependent control on the circadian clock, photomorphogenesis, and flowering time. Plant Cell 16:769–782.

Song, S.-J., Feng, Q.-N., Li, C.-L., Li, E., Liu, Q., Kang, H., Zhang, W., Zhang, Y., and Li, S. (2018). A tonoplast-associated calcium-signaling module dampens ABA signaling during stomatal movement. Plant Physiol. 177:1666.

Suarez-Lopez, P., Wheatley, K., Robson, F., Onouchi, H., Valverde, F., and Coupland, G. (2001). CONSTANS mediates between the circadian clock and the control of flowering in Arabidopsis. Nature 410:1116–1120.

Takeno, K. (2016). Stress-induced flowering: the third category of flowering response. J. Exp. Bot. 67:4925–4934.

Turck, F., Fornara, F., and Coupland, G. (2008). Regulation and identity of Florigen: FLOWERING LOCUS T moves center stage. Annu. Rev. Plant Biol. 59:573–594.

Valverde, F., Mouradov, A., Soppe, W., Ravenscroft, D., Samach, A., and Coupland, G. (2004). Photoreceptor regulation of CONSTANS protein in photoperiodic flowering. Science 303:1003–1006.

Wang, P., Zhao, Y., Li, Z., Hsu, C.-C., Liu, X., Fu, L., Hou, Y.-J., Du, Y., Xie, S., Zhang, C., et al. (2018). Reciprocal regulation of the TOR kinase and ABA receptor balances plant growth and stress response. Mol. Cell 69:100–112.e6.

Zhang, Y.-L., Li, E., Feng, Q.-N., Zhao, X.-Y., Ge, F.-R., Zhang, Y., and Li, S. (2015). Protein palmitoylation is critical for the polar growth of root hairs in Arabidopsis. BMC Plant Biol. 15:50.

Zhou, L.-Z., Li, S., Feng, Q.-N., Zhang, Y.-L., Zhao, X., Zeng, Y., Wang, H., Jiang, L., and Zhang, Y. (2013). Protein S-acyl transferase10 is critical for development and salt tolerance in Arabidopsis. Plant Cell 25:1093–1107.

Zoltowski, B. D., and Imaizumi, T. (2014). Structure and function of the ZTL/FKF1/LKP2 group proteins in Arabidopsis. The Enzymes 35:213–239.

