## Supplemental Methods and Figures for "A Calcium/Palmitoylation Switch Interfaces the Signaling Networks of Stress Response and Transition to Flowering"

### SUPPLEMENTAL MATERIALS AND METHODS

#### Hydroponics for salt tolerance phenotypes

Seeds were sterilized and vernalized for two days at 4°C, and then put on rockwool in hydroponic culture system with aerated nutrient solution (LAK medium, 1 mM $KH_2PO_4$, 2 mM $Ca(NO_3)_2$, 1 mM $MgSO_4$, 30 µM $H_3BO_3$, 10 µM $MnSO_4$, 1 µM $ZnSO_4$, 1 µM $CuSO_4$, 0.03 µM $(NH_4)_6Mo_7O_{24}$, and 100 µM $Fe^{2+}$ as *Sequestrene* 138-Fe, pH ~5.3) (Barragán et al., 2012) with or without 10 mM NaCl. Photographs were taken in 2 to 3 weeks after salt treatment.

#### Plasmid construction

For BiFC, full-length ORF sequences for *SOS2* and *SOS3* were amplified with indicated primers (Supplemental Table S1) to generate entry vectors with or without stop codons in the *pDONR*$^{TM}$*/Zeo* vector (Invitrogen, Carlsbad, CA, USA). cDNAs of *GI*, *SOS3* and *SOS3-1* were cloned under the *35S* promoter into vectors pDEST-$^{GW}$VYNE for *GI-VN,* and in pDEST-$^{GW}$VYCE for *SOS3-VC* or *SOS3-1-VC*, respectively, using Gateway cloning system (Gehl et al., 2009). The *SOS3-1* protein bears a three amino acid deletion in the third EF-hand motif and cannot bind calcium (Ishitani et al., 2000). The *GI* entry vector, *pENTR-1A-Amp-GI(s)* was given by Hong Gil Nam (DGIST, South Korea). All constructs used in this paper are in Supplemental Table S2.

The plasmids for transformation of GFP-translational fusions to SOS3, SOS3-G2A and SOS3-C3A in Arabidopsis were produced using the pGreen system (Hellens et al., 2000). Plasmid pGreenII 0000 was used as backbone to generate the final construct. A 0.67 kb *Eco*RV fragment containing the CaMV35S expression cassette was ligated in the plasmid pGreenII 0000 cut with *Pvu*II. A 1.18 kb fragment *Eco*RV fragment containing the *nos:hygR* marker cassette was ligated into pGreenII 0000 cut with *Hpa*I. The yeast expression vector, pYES2GFP was digested with *Hind*III and *Xba*I to isolate a 0.81 kb band containing the multiple cloning sites (MCS) and the GFP coding sequence. The *Xba*I site at the 3’ end of this fragment was filled with Klenow to create a blunt end, the purified MCS-GFP was ligated into the vector pGreenII 000 cut with *Hind*III and *Eco*RI (blunted) producing the final vector pGreenII 35S-GFP. The different alleles of SOS3 were generated by PCR using the forward primer: 5'-GGAAGCTTATGGGCTGCTCTGTATC (for SOS3 wild-type); 5'-GGAAGCTTATGGCCTGCTCTGTATC (for the Gly2 to Ala mutant); 5'-GGAAGCTTATGGGCGCCTCTGTATC (for the Cys3 to Ala mutant) and the reverse primer 5'- TCTGCGGCCGCGGAAGATACGTTTTGCAA using the SOS3 cDNA as template. *Hind*III and *Not*I restrictions sites are underlined in the primer sequence. The PCR fragments were digested with *Hind*III and *Not*I, purified and subcloned into the pGreenII vector cut with the same restriction enzymes.

For *sos3-1* complementation test, the different *SOS3* alleles were subcloned into the pBI321 vector (Martínez-Atienza et al., 2007). Plasmid *pYPGE-SOS3* (Guo et al., 2004) was digested with *Xba*I and *Xho*I to obtain a 0.67 kb fragment containing the wild-type SOS3 cDNA. Plasmid *pET-SOS3-G2A* (Ishitani et al., 2000) was digested with XbaI and XhoI to obtain a 0.69 kb fragment containing the mutant *SOS3-G2A* cDNA. The Cys3 to Ala (C3A) mutant of SOS3 was created by PCR using the forward primer: 5'-GATCTAGAATGGGCGCCTCTGTATC and the reverse primer 5'-AGCTCGAGGTTAGGAAGATACG. Primers add an *Xba*I site at the 5'end and an *Xho*I site at the 3' end of the amplified DNA. The *SOS3* alleles were ligated into the pBI321 vector cut with *Xba*I and *Xho*I.

The construct *proSOS3:SOS3-GFP* was used to mimic the expression of the native *SOS3* gene . For this, a translational fusion was constructed in which GFP was added to the C-terminus of a genomic copy of *SOS3*. The expression of SOS3 was driven by its own promoter (the 2kb promoter region upstream the ATG start codon). The *proSOS3:SOS3-GFP* construct was amplified and cloned into pGreenII as an *Xho*I-*Sph*I fragment. Forward and reverse oligonucleotides for cloning were SOS3 promoter: 5-CGGCATGCAGATCTAAAAACAGGTAATGAGAATTTGG-3´ and Fusion SOS3-GFP: 5´-

CCACCTCGAGCGGAAGATACGTT-3´. Transgenic plants in the *sos3-1* background were selected using 15 mg/l of Hygromycin in ½ MS plates. Homozygous plants from T2 generation were for tested for complementation of salt-sensitivity and used for further experimentation.

To test fatty acid modification of SOS3 in yeast cells, the wild-type allele of *SOS3*, the non-myristoylatable mutant SOS3-G2A and the non-palmitoylatable mutant SOS3-C3A were generated by PCR using the following forward primers: 5'-GATCTAGAATGGGCTGCTCTGTATC (for SOS3 wild-type); 5'-GATCTAGAATGGCCTGCTCTGTATC (for the Gly2 to Ala mutant; G2A); 5'-GATCTAGAATGGGCGCCTCTGTATC (for the Cys3 to Ala mutant, C3A) and the reverse primer 5'- TCTGCGGCCGCGGAAGATACGTTTTGCAA. *Xba*I and *Not*I restrictions sites are underlined in the primer sequence. PCR fragments were digested with *Xba*I and *Not*I, purified and subcloned into a pYPGE15 vector (Brunelli and Pall, 1993) modified to include the 6xHis-tag.

For SOS3 S-acylation test using transient tobacco expression and acyl resin-assisted capture, the wild-type allele of SOS3, the non-myristoylatable mutant SOS3-G2A, the non-palmitoylatable mutant SOS3-C3A and the non-myristoylatable, non-palmitoylatable double mutant SOS3-G2A/C3A were generated by PCR using the following forward primers: 5´-AAAAAGCAGGCTTCATGGGCTGCTCTGTATCGAAGAAG (for SOS3 WT); 5´-AAAAAGCAGGCTTCATGGCATGCTCTGTATCGAAGAAGAAG (for the G2A mutant); 5´-AAAAAGCAGGCTTCATGGGCGCATCTGTATCGAAGAAGAAG (for the C3A mutant); 5´-AAAAAGCAGGCTTCATGGCAGCATCTGTATCG AAGAAGAAGAAG (for the G2A and C3A double mutant), and the reverse primer 5´-AGAAAGCTGGGTCGGAAGATACGTTTTGCAATTCCATTTCT for all the above PCRs. Amplification products were subcloned using LR II clonase (Invitrogen) into the Gateway-compatible pYL436 vector, which confers C-terminal TAP tags (Rubio et al., 2005).

**Transient expression in tobacco leaves**

*Agrobacterium tumefaciens* strain GV 3101 was transformed with constructs indicated in figure legends. *Agrobacterium* grew in LB media supplemented with 10 mM MES, 20 µM acetosyringone, and antibiotics dependent on the constructs and culture media were washed with infiltration solution (10 mM $MgCl_2$, 10 mM MES, and 100 µM acetosyringone). *Agrobacterium* transformed with P19, repressor of silencing was included (Lakatos et al., 2004). For co-infiltration, each of *Agrobacterium* cultures was $OD_{600}$0.5 in the final infiltration solution, which was infiltrated to leaves of three to four-week old tobacco plants (*Nicotiana benthamiana)* through stomata. Infiltrated tobacco plants were incubated 2-3 days, and then treated if needed with 100 mM NaCl or 3 mM $CaCl_2$ solutions by infiltration into leaves in the morning (ZT2). After 10 h, salt-treated tobacco leaves were harvested for further experiments such as co-IP or confocal laser fluorescence microscopy.

**2-Bromo-Palmitate treatment and spinning-disc confocal microscopy**

Plants were sown in ½ MS plates with 1% sucrose in a long-day chamber at 21°C. Five days after germination the plants were incubated in liquid in ½ MS with 1% sucrose media, with or without 50 µM 2-bromopalmitate (2-BrP; Sigma-Aldrich), and with or without 100 mM NaCl for 24 h keeping the same growth conditions. For controls, a mock treatment with the same volume of ethanol (2-BrP solvent) was added to samples without 2-BrP.

For regular confocal microscopy, plants were incubated for 10 min with 0.1% Triton and 0.2 µg/ml DAPI (Sigma-Aldrich), washed three times with ½ MS media and then images were taken. DAPI was exited with 405 laser and emission collected between 440-480 nm. GFP was exited with 488 laser and emission collected between 490-550 nm. Arabidopsis seedlings were imaged between slide and cover glass. The pictures were taken using a Zeiss LSM 780 with the objective 40x 1.1NA Water LD C-Apochromat Korr M27 (DIC). Image acquisitions were performed sequentially and analyzed using FIJI software, version 1.57. DAPI signal was used as a ROI to establish the localization of the nuclei to measure the fluorescence intensity (mean gray value (a.u.)) of *proSOS3::SOS3GFP* plants in different cellular compartments.

For spinning-disc confocal microscopy, pictures were generated using a confocal microscope equipped with a 113 CSU-W1 spinning disc head (Yokogawa, Tokyo, japan) fitted to a Nikon Eclipse Ti-E-inverted microscope with a CFI PlanApo VC 60x N.A. 1.40 oil immersion objective and an EM-CCD ImageEM 1K (c9100-14) camera. GFP was imaged using a 488 nm solid-state diode laser and a 525/50 nm emission filter. The fluorescence intensity (mean grey value (a.u.)) of *proSOS3::SOS3GFP* plants was measured in different cellular compartments. Image processing was done with Fiji software, version 1.57.

**Immunoblotting and immunoprecipitation**

Total proteins were obtained in extraction buffer (100 mM Tris-HCl, pH 7.5, 150 mM NaCl, 0.5% NP-40, 1 mM EDTA, 3 mM DTT, and protease inhibitors 1 mM PMSF, 5 µg/ml leupeptin, 1 µg/ml aprotinin, 1 µg/ml pepstatin, 5 µg/ml antipain, 5 µg/ml chymostatin, 2 mM sodium orthovanadate ($Na_3VO_4$), 2 mM sodium fluoride (NaF) and 50 µM MG132) (Kim et al., 2013). For Arabidopsis protein extraction, 1% NP-40 was added to the extraction solution.

**Chromatin immunoprecipitation (ChIP) assay**

Procedures of fixation and isolation of chromatin were performed as described (Sawa et al., 2007; Saleh et al., 2008). Two-week old Arabidopsis seedlings (*GI-GFPox* and *SOS3-GFPox*) treated with 100 mM NaCl for 10 h, and then fixed with 1% formaldehyde under vacuum for 15 min. Subsequently 0.125 M glycine was added to stop cross-linking. Nuclei were isolated in nuclei isolation buffer (0.25 M sucrose, 15 mM PIPES pH 6.8, 5 mM $MgCl_2$, 60 mM KCl, 15 mM NaCl, 1 mM $CaCl_2$, 0.9% Triton X-100, 1 mM PMSF, 2 µg/ml pepstatin A, and 2 µg/ml aprotinin), lysed in ice-cold nuclei lysis buffer (50 mM HEPES pH 7.5, 150 mM NaCl, 1mM EDTA, 1% SDS, 0.1% sodium deoxycholate, and 1% Triton X-100, 1 µg/ml pepstatin A, and 1 µg/ml aprotinin), and sonicated to shear DNA to approximately 500 bp. The chromatin samples were diluted 10-fold in lysis buffer and incubated with salmon sperm

DNA/protein A agarose bead (Millipore, Darmstadt, Germany) for 1 h. Appropriate antibody (α-GFP, Abcam, #ab290) was added to the supernatants and incubated overnight. The beads were washed with ice cold low salt wash buffer (150 mM NaCl, 20 mM Tris-HCl, pH8, 0.2% SDS, 0.5% Triton X-100, and 2 mM EDTA), high salt wash buffer (500 mM NaCl, 20 mM Tris-HCl, pH8, 0.2 % SDS, 0.5% Triton X-100, and 2 mM EDTA), LiCl wash buffer (0.25 M LiCl, 1% sodium deoxycholate, 10 mM Tri-HCl pH8, 1% NP-40, and 1 mM EDTA) and TE buffer (1 mM EDTA and 10 mM Tris-HCl pH8) sequentially. Immunocomplexes were eluted with elution buffer (0.5% SDS and 0.1 M $NaHCO_3$) at room temperature. Reverse-crosslinking was done by incubating with NaCl overnight at 65°C and subsequent incubation with proteinase K. DNA was purified by phenol-chloroform extraction and rescued by ethanol precipitation with 20 µg glycogen (Fermentas). DNA was resuspended in 50 µl TE buffer and 2 µl aliquots were used for qPCR. The primers used in this experiment are in Supplemental Table S2.

**Detection of SOS3 S-acylation by differential alkylation**

Wild-type SOS3 and mutant proteins G2A, C3A and G2A/C3A were expressed in yeast grown overnight in YPD medium at 30°C. Cells were collected by centrifugation and resuspended in buffer A (50 mM $NaH_2PO_4$, pH 8.0, 300 mM NaCl). The cell suspension was lysed using glass beads following standard protocols. Unbroken cells and debris were discarded after centrifugation at 3000 g for 10 min. The supernatant was centrifuged at 20,000 g for 30 min to obtain a clarified total protein extract. Proteins were treated with 30 mM N-ethylmaleimide (NEM) to block free cysteine thiols, precipitated with 10% trichloroacetic acid (TCA), and resuspended in a denaturing buffer consisting of 50 mM Tris-HCl (pH 7.8), 4 M urea, 2% SDS and 2.5% glycerol. Thereafter samples were incubated in the presence or absence of 0.7 M hydroxylamine for 1 h at 25°C to break palmitoyl-thioester bonds. Excess reagent was removed by TCA precipitation and samples were resuspended in a denaturation buffer containing 10 mM methyl-$PEG_{24}$-maleimide, $MM(PEG)_{24}$, which alkylates newly formed cysteine thiols. Proteins were resolved in 11% acrylamide SDS-

PAGE gels and subjected to western blot analysis using α-SOS3 antibodies (Ishitani et al., 2000) at 1:2000 dilution.

**Analysis of SOS3 *S*-acylation *in planta* through acyl resin-assisted capture (acyl-RAC)**

*Agrobacterium* cells transformed with the constructs of wild-type SOS3 and mutant SOS3 C-terminally tagged with TAP tag were infiltrated in *N. benthamiana* leaves, and wild-type and mutant SOS3 proteins were transiently expressed during 72 h. For each sample 1 g of leaf tissue was ground in liquid nitrogen and resuspended in 7.5 ml buffer containing 100 mM Tris-HCl (pH 7.5), 1% SDS, 1 mM PMSF, 1 mM EDTA and 30 mM N-ethylmaleimide. Leaf suspensions were first heated at 40°C for 10 min and then mixed gently for 1 h at 23°C. Debris was removed by centrifugation at 12000 *g* for 10 min at 23°C and supernatants were passed through 0.2 µm filters before precipitation with 85% acetone at -20°C for 1 h. Precipitated proteins were collected by centrifugation at 12000 *g* for 10 min at 4ºC. Pellets were washed once in 70% acetone and dried at 42°C for 30 min. Each protein pellet was dissolved in 20 ml binding-buffer consisting of 100 mM Tris-HCl (pH 7.5), 1% SDS, 1 mM EDTA and 4 M urea. Solutions were cleared by centrifugation at 12000 *g* for 10 min at 23°C and filtered through 0.2 µm filters. Fifty-µl aliquots were withdrawn from these solutions to be analyzed by western blot as "input". Sample solutions were divided in two 10-ml parts and hydroxylamine was added at 0.5 M final concentrations to one of these parts, whereas the other served as untreated control. Each sample solution was incubated with 200 µl thiopropyl-sepharose 6B resin (Sigma) overnight at 23°C under gentle mixing. The resin slurries were washed with 5 ml binding-buffer and covalently bound proteins were eluted in 200 µl of a buffer containing 20 mM Tris-HCl (pH 6.8), 0.4% SDS and 50 mM DTT.

**RNA isolation and Q-RT PCR**

Total RNA (3 µg) extracted using RNeasy Plant Mini Kit (Qiagen, Hilden, Germany) and treated with DNase (Sigma) was used for synthesis of first-strand cDNA using first-strand cDNA using the ReverTra Ace-α-® (Toyobo Co. Ltd, Osaka, Japan). Amplified products were detected using iQ™ SYBR® Green Supermix (Bio-Rad) in a thermal cycler (CFX384 C1000TM Real time system. Bio-Rad). The efficiency value of amplification for each primer set was checked by measuring the abundance of transcripts from cDNA dilutions according to the manufacture guide book (real-time PCR applications guide, Bio-Rad). Q-RT–PCR conditions for *CO* and *FT* were as follows: 95°C for 5 min, 55 cycles of 95°C for 15 s, 55°C for 15 s and 72°C for15 s, followed by 95°C for 10 s, 65°C for 5 s, and 95°C for 5 s for melting curves. Each data point shown is the average of three independent amplifications and each biological replica had three technical replicas. Primers used for Q-RT PCR are in Supplemental Table S1.

**Supplemental Table S1. List of primers.**

| Primers | Sequence | Reference | Purpose |
|---|---|---|---|
| attB1 sequence | AAAAAGCAGGCTNN | Invitrogen | Gateway |
| attB2 sequence | AGAAAGCTGGGTN | Invitrogen | |
| attB1 SOS3-F | AAAAAGCAGGCTTCATGGGCTGCTCTGTATCGAA | This study | Plasmid construction |
| attB1SOS3-G2A-F | AAAAAGCAGGCTTCATGGCCTGCTCTGTATCG | This study | |
| attB1SOS3-C3A-F | AAAAAGCAGGCTTCATGGGCGCCTCTGTATCGAAG A | This study | |
| attB1-SOS3-K/A F | AAAAAGCAGGCTTCATGGGCTGCTCTGTATCG | This study | |
| attB2-SOS3-R/No Stop | AGAAAGCTGGGTCGGAAGATACGTTTTGCAAT | This study | |
| attB1 SOS2-F | AAAAAGCAGGCTTCATGACAAAGAAAATGAGAAG | Kim et al., 2013 | |
| attB2-SOS2-R/Stop | AGAAAGCTGGGTTCAAAACGTGATTGTTCTGAG | Kim et al., 2013 | |
| attB2-SOS2-R/No Stop | AGAAAGCTGGGTCAAACGTGATTGTTCTGAGAAT | Kim et al., 2013 | |
| attB1-FKF1-F | AAAAAGCAGGCTTCATGGCGAGAGAACATGCGAT | This study | |
| attB2-FKF1-R/Stop | AGAAAGCTGGGTCTTACAGATCCGAGTCTTGCC | This study | |
| FwPr-SOS3-GFP | CGGCATGCAGATCTAAAAAACAGGTAATGAGAATTT GG | This study | |
| RvPr-SOS3-GFP | CCACCTCGAGCGGAAGATACGTT | This study | |
| UBQ-qRT-F | GACGCTTCATCTCGTCC | This study | Real-time PCR |
| UBQ-qRT-R | GTAAACGTAGGTGAGTCC | This study | |
| CO-qRT-F | ATTCTGCAAACCCACTTGCT | Kim et al., 2013 | |
| CO-qRT-R | CCTCCTTGGCATCCTTATCA | Kim et al., 2013 | |
| FT-qRT-F | CTGGAACAACCTTTGGCAAT | Kim et al., 2013 | |
| FT-qRT-R | AGCCACTCTCCCTCTGACAA | Kim et al., 2013 | |
| CO_pro_amp_#4(A)_ LP | TATGGTCCCTCGACTCTTATTCTCT | Sawa et al., 2007 | ChIP |
| CO_pro_amp_#4(A)_ RP | GCCTTCGGATAACTGTTACGAGTAA | Sawa et al., 2007 | |
| CO_pro_amp_#5(B)_ LP | ATTTACTCTTCATGAACTCGAACCA | Sawa et al., 2007 | |
| CO_pro_amp_#5(B)_ RP | ACTGGTTTTACGATGAATGTAATGG | Sawa et al., 2007 | |
| CO_pro_amp_#8(C)_ LP | TGGTTACCAAGTGCAAATTTCTACA | Sawa et al., 2007 | |
| CO_pro_amp_#8(C)_ RP | GAGAATCATATCGGAAAAGTGACATGAA | Sawa et al., 2007 | |
| UBQ10_For_LP | TCCAGGACAAGGAGGTATTCCTCCG | Sawa et al., 2007 | |
| UBQ10_Rev_RP | CCACCAAAGTTTTACATGAAACGAA | Sawa et al., 2007 | |

**Supplemental Table S2. List of constructs**.

| Name | Construct | Vector name | Reference | Purpose |
|---|---|---|---|---|
| GI-VN | *35S:GI-VN* | pDEST-$^{GW}$VYNE | Kim et al., 2013 | BIFC |
| VC-SOS2 | *35S:VC-SOS2* | pDEST-VYCE(R)$^{GW}$ | Kim et al., 2013 | |
| SOS3-VC | *35S: SOS3-VC* | pDEST- $^{GW}$VYCE | This study | |
| SOS3-1-VC | *35S:SOS3-1-VC* | pDEST-$^{GW}$VYCE | This study | |
| SOS3-G2A-VC | *35S:SOS3-G2A-VC* | pDEST- $^{GW}$VYCE | This study | |
| SOS3-C3A-VC | *35S:SOS3-C3A-VC* | pDEST- $^{GW}$VYCE | This study | |
| VC-FKF1 | *35S:VC-FKF1* | pDEST-VYCE(R)$^{GW}$ | This study | |
| GI-HA | *35S:GI-HA* | pGWB14 | Kim et al., 2013 | Transient expression |
| GI-GFP | *35S:GI-GFP* | pK7WGF | Kim et al., 2013 | |
| SOS3-MYC | *35S:SOS3-MYC* | pCambia1300PT | Kim et al., 2013 | |
| SOS3-FLAG | *35S:SOS3-FLAG* | pGWB11 | This study | |
| SOS3-GFP | *35S:SOS3-GFP* | Modified pGreen0000 | This study | |
| SOS3-G2A-GFP | *35S:SOS3-G2A-GFP* | Modified pGreen0000 | This study | |
| SOS3-C3A-GFP | *35S:SOS3-C3A-GFP* | Modified pGreen0000 | This study | |
| MYC-SOS3-1 | *35S:MYC-SOS3-1* | pEarleygate203 | This study | |
| MYC-FKF1 | *35S:MYC-FKF1* | pEarleygate203 | This study | |
| SOS3-TAP | *2x35S:SOS3-TAP* | pYL436 | This study | |
| SOS3-G2A-TAP | *2x35S:SOS3-G2A-TAP* | pYL436 | This study | |
| SOS3-C3A-TAP | *2x35S:SOS3-C3A-TAP* | pYL436 | This study | |
| SOS3-G2AC3A-TAP | *2x35S:SOS3-G2A C3A-TAP* | pYL436 | This study | |
| SOS3 | *35S:SOS3* | pBI321 | This study | Transgenic plants |
| SOS3-G2A | *35S:SOS3-G2A* | pBI321 | This study | |
| SOS3-C3A | *35S:SOS3-C3A* | pBI321 | This study | |
| proSOS3:SOS3-GFP | *proSOS3:SOS3-GFP* | pGreenII | This study | |
| SOS3-H6x | *PGK1:SOS3-H6x* | pYPGE15 | This study | Expression in yeast |
| SOS3-G2A-H6x | *PGK1:SOS3-G2A-H6x* | pYPGE15 | This study | |
| SOS3-C3A-H6x | *PGK1:SOS3-C3A-H6x* | pYPGE15 | This study | |
| SOS3-G2A/C3A-H6x | *PGK1:SOS3-G2A/C3A-H6x* | pYPGE15 | This study | |

**Supplemental References**


Barragán, V., Leidi, E. O., Andrés, Z., Rubio, L., De Luca, A., Fernández, J. A., Cubero, B., and Pardo, J. M.(2012). Ion exchangers NHX1 and NHX2 mediate active potassium uptake into vacuoles to regulate cell turgor and stomatal function in Arabidopsis. *Plant Cell* 24:1127–1142.

Brunelli, J. P., and Pall, M. L. (1993). A series of yeast shuttle vectors for expression of cDNAs and other DNA sequences. *Yeast* 9:1299–1308.

Gehl, C., Waadt, R., Kudla, J., Mendel, R.-R., and Hänsch, R. (2009). New GATEWAY vectors for high throughput analyses of protein–protein interactions by bimolecular fluorescence complementation. *Mol. Plant* 2:1051–1058.

Guo, Y., Qiu, Q.-S., Quintero, F. J., Pardo, J. M., Ohta, M., Zhang, C., Schumaker, K. S., and Zhu, J.-K. (2004). Transgenic evaluation of activated mutant alleles of SOS2 reveals a critical requirement for its kinase activity and C-terminal regulatory domain for salt tolerance in *Arabidopsis thaliana*. *Plant Cell* 16:435–449

Hellens, R. P., Edwards, E. A., Leyland, N. R., Bean, S., and Mullineaux, P. M. (2000). pGreen: a versatile and flexible binary Ti vector for Agrobacterium-mediated plant transformation. *Plant Mol. Biol.* 42:819–832

Ishitani, M., Liu, J., Halfter, U., Kim, C.-S., Shi, W., and Zhu, J.-K. (2000). SOS3 function in plant salt tolerance requires N-myristoylation and calcium binding. *Plant Cell* 12:1667–1677.

Kim, W.-Y., Ali, Z., Park, H. J., Park, S. J., Cha, J.-Y., Perez-Hormaeche, J., Quintero, F. J., Shin, G., Kim, M. R., Qiang, Z., et al. (2013). Release of SOS2 kinase from sequestration with GIGANTEA determines salt tolerance in Arabidopsis. *Nat. Commun.* 4:1352.

Lakatos, L., Szittya, G., Silhavy, D., and Burgyán, J. (2004). Molecular mechanism of RNA silencing suppression mediated by p19 protein of tombusviruses. *EMBO J.* 23:876–884.

Martínez-Atienza, J., Jiang, X., Garciadeblas, B., Mendoza, I., Zhu, J.-K., Pardo, J. M., and Quintero, F. J. (2007). Conservation of the salt overly sensitive pathway in rice. *Plant Physiol.* 143:1001–1012.

Rubio, V., Shen, Y., Saijo, Y., Liu, Y., Gusmaroli, G., Dinesh-Kumar, S. P., and Deng, X. W. (2005). An alternative tandem affinity purification strategy applied to Arabidopsis protein complex isolation: Alternative tandem affinity purification in Arabidopsis. *Plant J.* 41:767–778.

Saleh, A., Alvarez-Venegas, R., and Avramova, Z. (2008). An efficient chromatin immunoprecipitation (ChIP) protocol for studying histone modifications in Arabidopsis plants. *Nat. Protoc.* 3:1018–1025

Sawa, M., Nusinow, D. A., Kay, S. A., and Imaizumi, T. (2007). FKF1 and GIGANTEA complex formation is required for day-length measurement in Arabidopsis. *Science* 318:261–265.

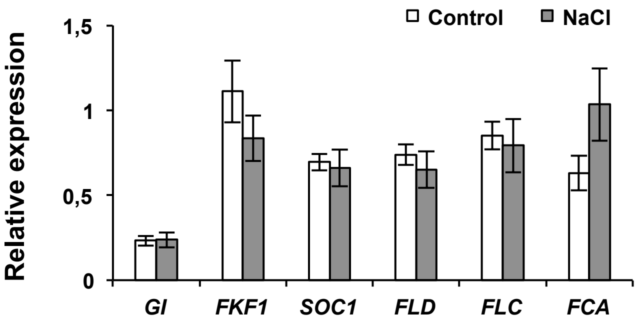


**Figure S1. Expression under salt stress of genes regulating flowering.**

Two-week old wild-type plants (Col-*gl1*) grown in long-day were treated with 100 mM NaCl at ZT0. Transcript levels of *GI, FKF1, FLD*, *SOC1*, *FLC* and *FCA* at ZT16 were measured by qRT-PCR. Bars represent means ± SEM from three technical replicates. Means of control and treated plants for each gene were not significantly different by Student's t-test at $p>0.1$. The experiment was repeated at least twice with similar results.

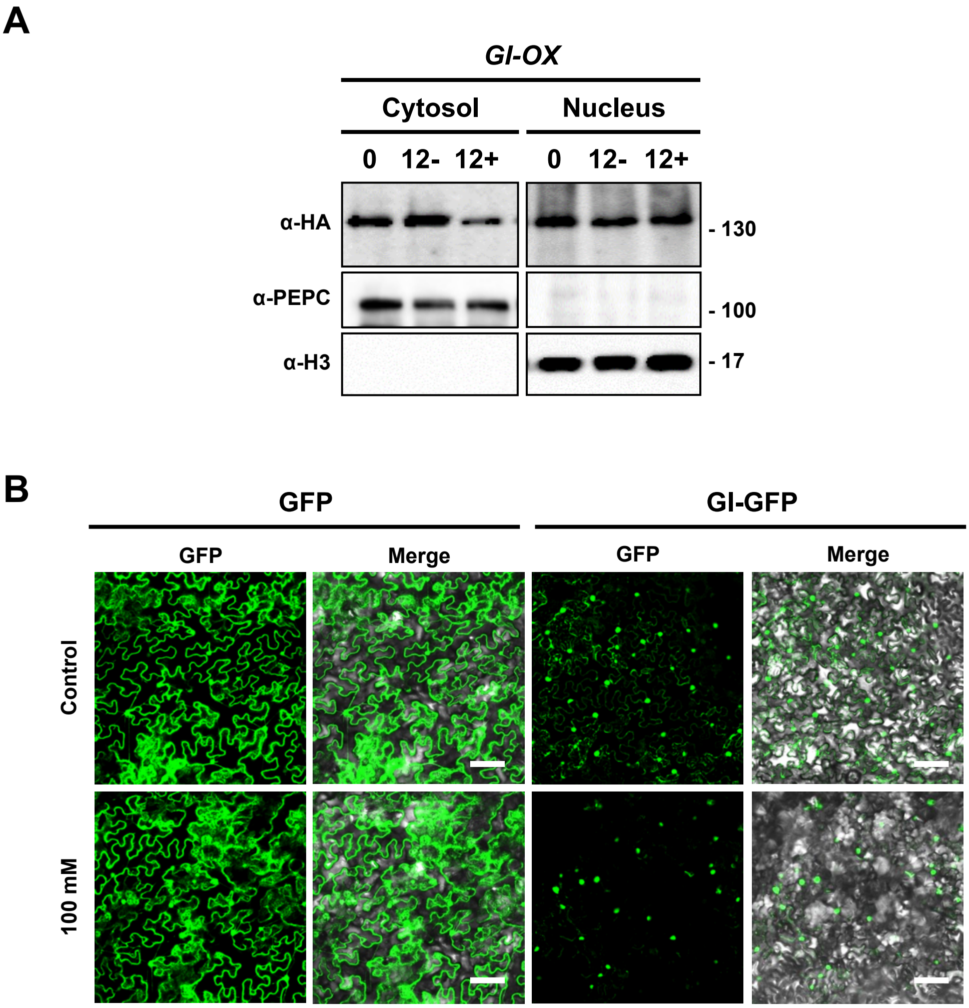


**Figure S2. Salt induced degradation of GI protein occurs in cytosol.**

(**A**) Two-week old *GI-OX* plants were treated with (12+) or without (12-) 100 mM NaCl for 12 h. Cytosolic and nuclear proteins were extracted and submitted to western blotting. α-PEPC and α-H3 antibodies were used for cytosolic and nuclear markers, respectively.

(**B**) Tobacco leaves transiently expressing *GI-GFP* were treated with or without 100 mM NaCl for 4 h. GFP signals were detected under confocal microscope. Bar represents 100 μm.

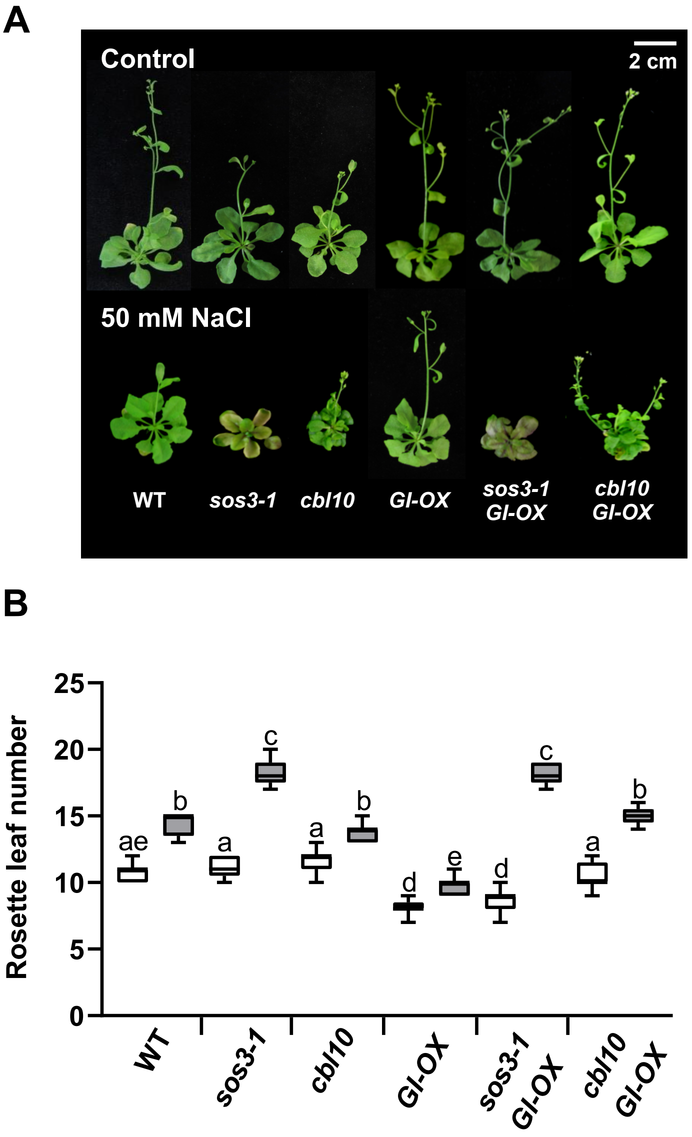


**Figure S3. CBL10 is not involved in the salt-induced late flowering.**

(**A**) Eight-day old plants of Col-0 (WT), *sos3-1, cbl10,* and the same genotypes over-expressing *GI* (*GI-OX, sos3-1 GI-OX, cbl10 GI-OX*) were transferred to MS media supplemented with 50 mM NaCl. Rosette leaf number was counted at bolting as flowering time. (**B**) Box plots of flowering time. Center lines show the medians; box limits indicate the 25th and 75th percentiles; whiskers extend to the minimum and maximum (n=9). Letters indicate significantly different means at $p<0.01$, Fisher's LSD test. (**C**) Quantitation of BiFC in *Nicotiana* leaves expressing SOS3 and GI. Leaves were treated for 9 h with 100 mM NaCl, or 3 mM $CaCl_2$, with or without 2 mM of EGTA. Water was used as mock treatment and the SOS3-1 mutant protein as the negative control. The fluorescence intensity was measured in frames of 0.4 $mm^2$ captured with x20 magnification objective. Data is shown as box plots, as in (**B**). Letters indicate significantly different means, $p < 0.05$ by Fisher's LSD test, $n \geq 3$.

A

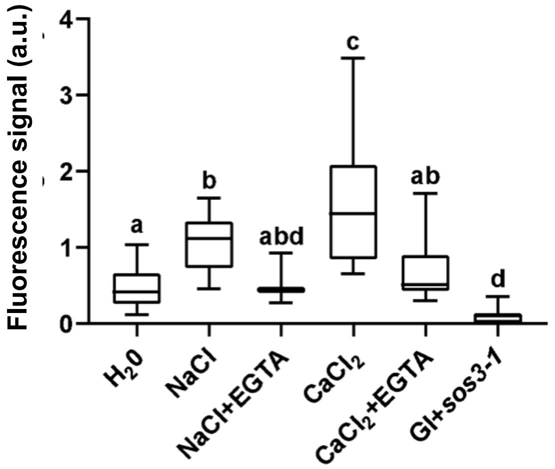


B

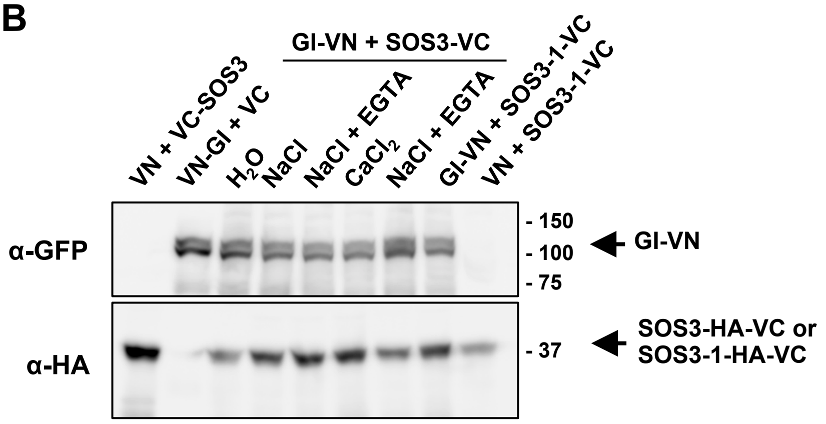


**Figure S4. BiFC of SOS3 and GI in *Nicotiana* leaves.**

(**A**) Quantitation of BiFC shown in Figure 3D**.** Leaves were treated for 9 h with 100 mM NaCl, or 3 mM $CaCl_2$, with or without 2 mM of EGTA. Water was used as mock treatment and the SOS3-1 mutant protein as the negative control. The fluorescence intensity was measured in frames of 0.4 $mm^2$ captured with x20 magnification objective. Data is shown as box plots. Letters indicate significantly different means, $p < 0.05$ by Fisher's LSD test, n ≥ 3. (B) Expression and stability of proteins used for BiFC analyses. Western blot with total proteins from the tobacco leaves used for the BiFC assay shown in Figure 3D.

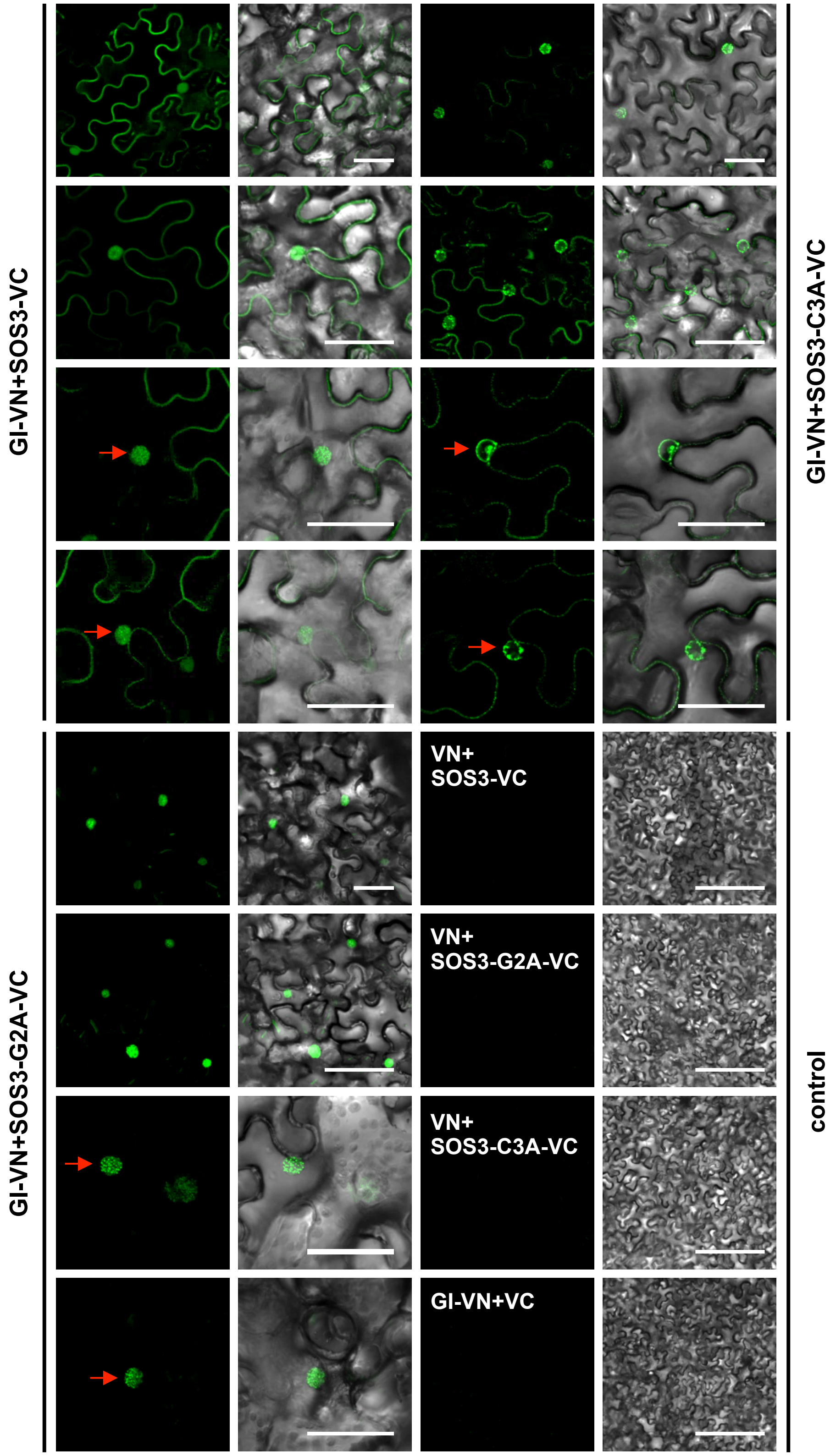


**Figure S5. Non-palmitoylated SOS3 fails to interact with GI in the nucleus.**
Images complement those in Figure 5D. Tobacco leaves were infiltrated with constructs as indicated. Fluorescent signals of GI and SOS3 interaction were detected under confocal microscope. Empty vectors were used as controls as indicated. Bars represent 50 μm.

A

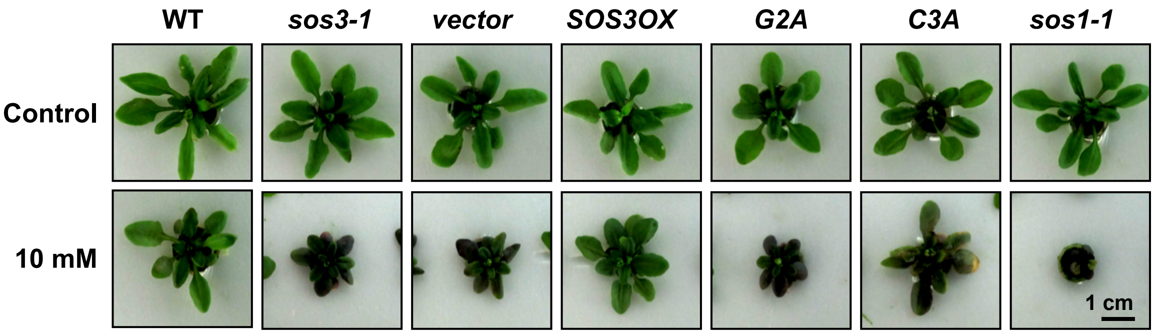


B

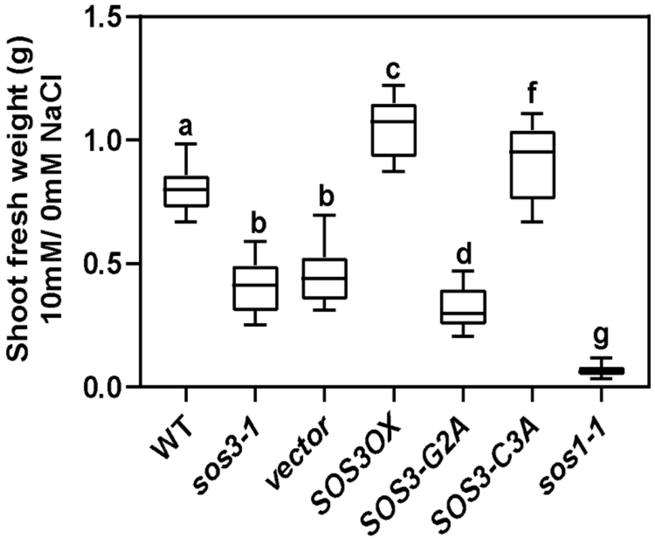


**Figure S6. Non-palmitoylated SOS3-C3A can partially rescue the salt sensitivity of *sos3-1.***
(**A**) Col-0 (WT), *sos3-1*, and the *sos3-1* transformed with an empty vector (vector) or over-expressing the wild-type SOS3 protein (SOS3OX), the non-myristoylated (SOS3-G2A) and the non-palmitoylated (SOS3-C3A) mutant proteins were grown in hydroponics with nutrient media supplemented or not with 10 mM NaCl. The *sos1-1* was used as a salt-hypersensitive control. Photographs were taken after 2 weeks of salt treatment.
(**B**) Shoot fresh weight of plants shown in (**A**) was measured and normalized to that of non- treated plants. Shown are the Box plots: center lines show the medians; box limits indicate the 25th and 75th percentiles; whiskers extend to the minimum and maximum (n≥9). Means followed by the same alphabet letter are not significantly different at $p<0.05$ by the Fisher's LSD.

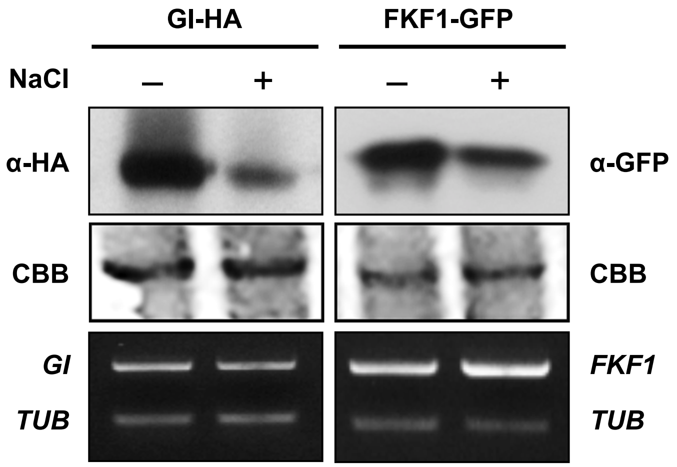


**Figure S7. FKF1 protein abundance upon exposure to salt**

*GI* and *FKF1* overexpressing plants carrying *35S:GIHA* (*GI-HA*) and *35S:FKF1-GFP* (*FKF1-GFP*), respectively, were treated with 100 mM NaCl at ZT0. GI and FKF1 protein abundance was evaluated after 12 h NaCl treatment by western blotting with antibodies α-HA and α-GFP (upper panel). Coomassie brilliant blue (CBB)-stained blots are shown as loading control (middle). Total RNA was extracted from *GI-HA* and *FKF1-GFP* plants and submitted to cDNA synthesis. *GI*, *FKF1*, and *Tubulin* (*TUB*) transcript levels were evaluated by RT-PCR (bottom).

A

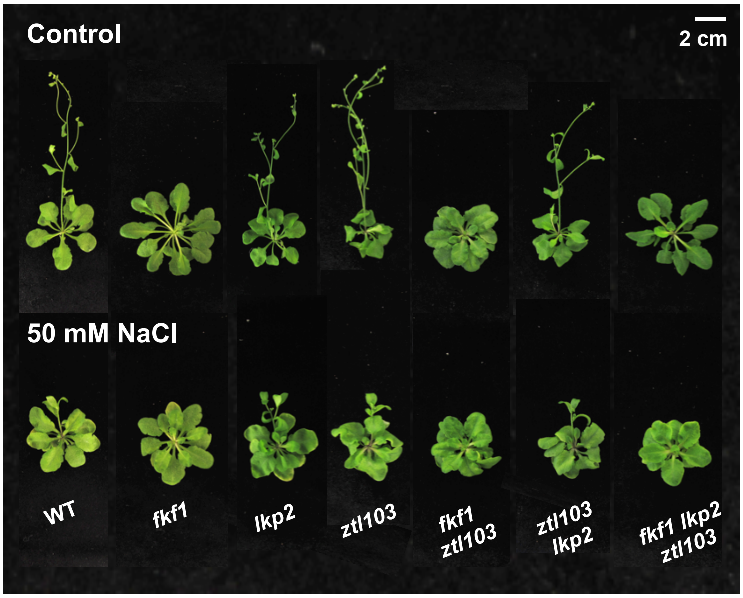


B

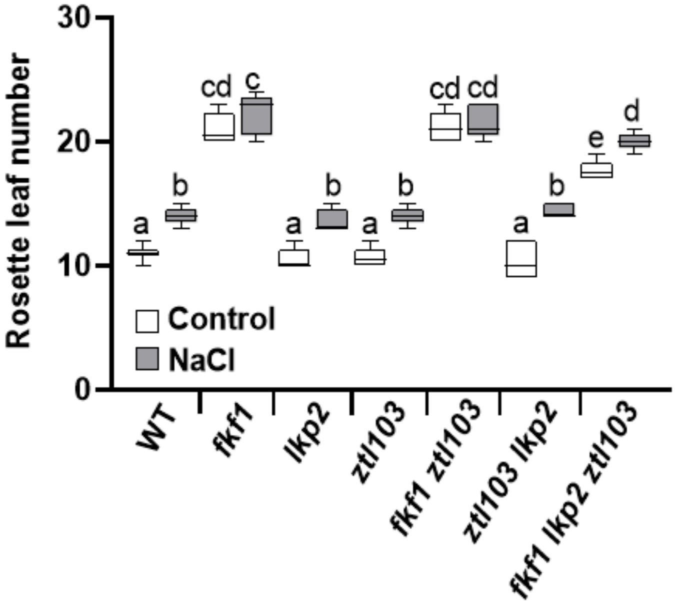


**Figure S8. Flowering time of blue-light photoreceptor mutants in saline media.**
(**A**) Eight-day-old plants including wild-type, single mutants *fkf1, lkp2*, *ztl103,* and their double and triple mutants were transferred to 50 mM NaCl. The picture was taken at bolting and rosette leaf numbers were counted as flowering time. (**B**) Box plots of flowering time. Center lines show the medians; box limits indicate the 25th and 75th percentiles; whiskers extend to the minimum and maximum (n≥5). Letters indicate statistically difference means at $p < 0.01$ by the Fisher's LSD; means with the same letter are statistically similar.

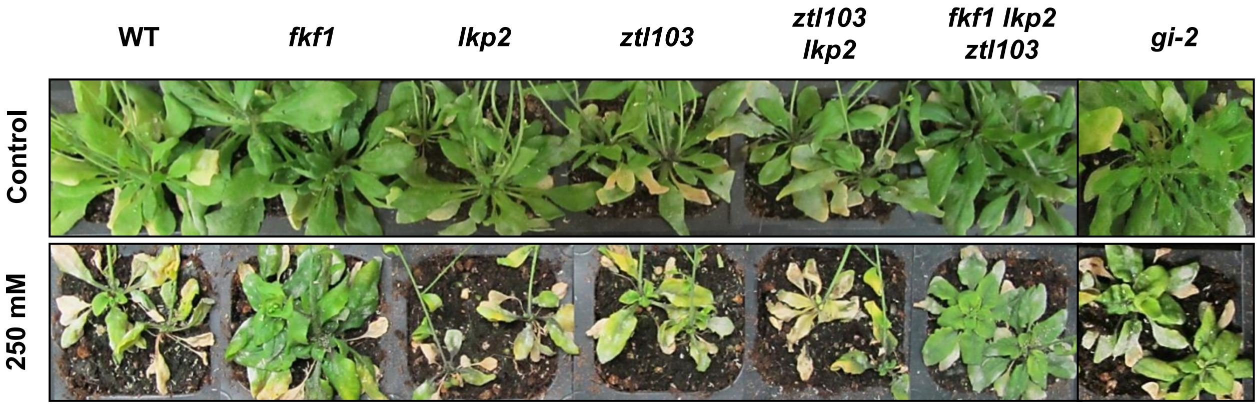


**Figure S9. The *fkf1* mutants exhibit increased salt tolerance, but not *lkp2* and *ztl103*.**
Col-0 (WT), *fkf1*, *lkp2*, *ztl103*, *ztl103 lkp2, fkf1 lkp2 ztl103* and *gi-2* plants were grown on soil for 3 weeks, and then treated or not with 250 mM NaCl for 2 weeks. Plants shown are representative of the 15 individual plants that were examined for each line.

A

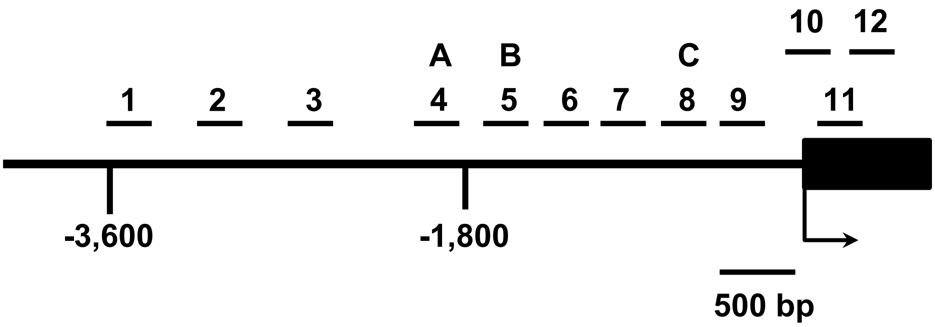


B

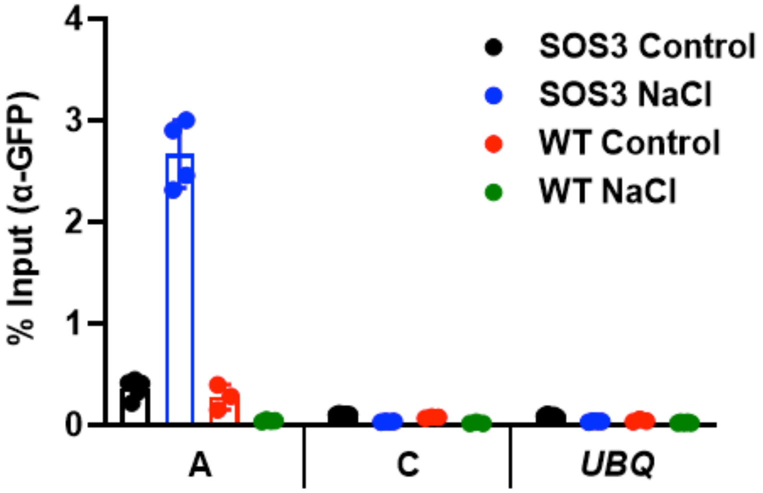


**Figure S10. SOS3 associates with the *CO* promoter.**

(**A**) Schematic drawing of the *CO* promoter and the amplicon locations (A, B, and C) for ChIP analysis, as defined by Sawa et al. (2007). The amplicon regions 4 and 8 in Sawa et al. (2007) are used as A-B and C, respectively, in this study.

(**B**) Chromatin isolated from two-week-old *SOS3-GFPox* (SOS3) and wild-type plants (WT) treated with or without 100 mM NaCl for 10 hours was immunoprecipitated with an α-GFP antibody. Immunoprecipitated and input DNA were used as templates for qPCR using primers specifically targeting to the amplicons A, C; *UBQ10* was used as control. Data is fragment enrichment as percent of input DNA. Error bar represents SEM (n≥3).

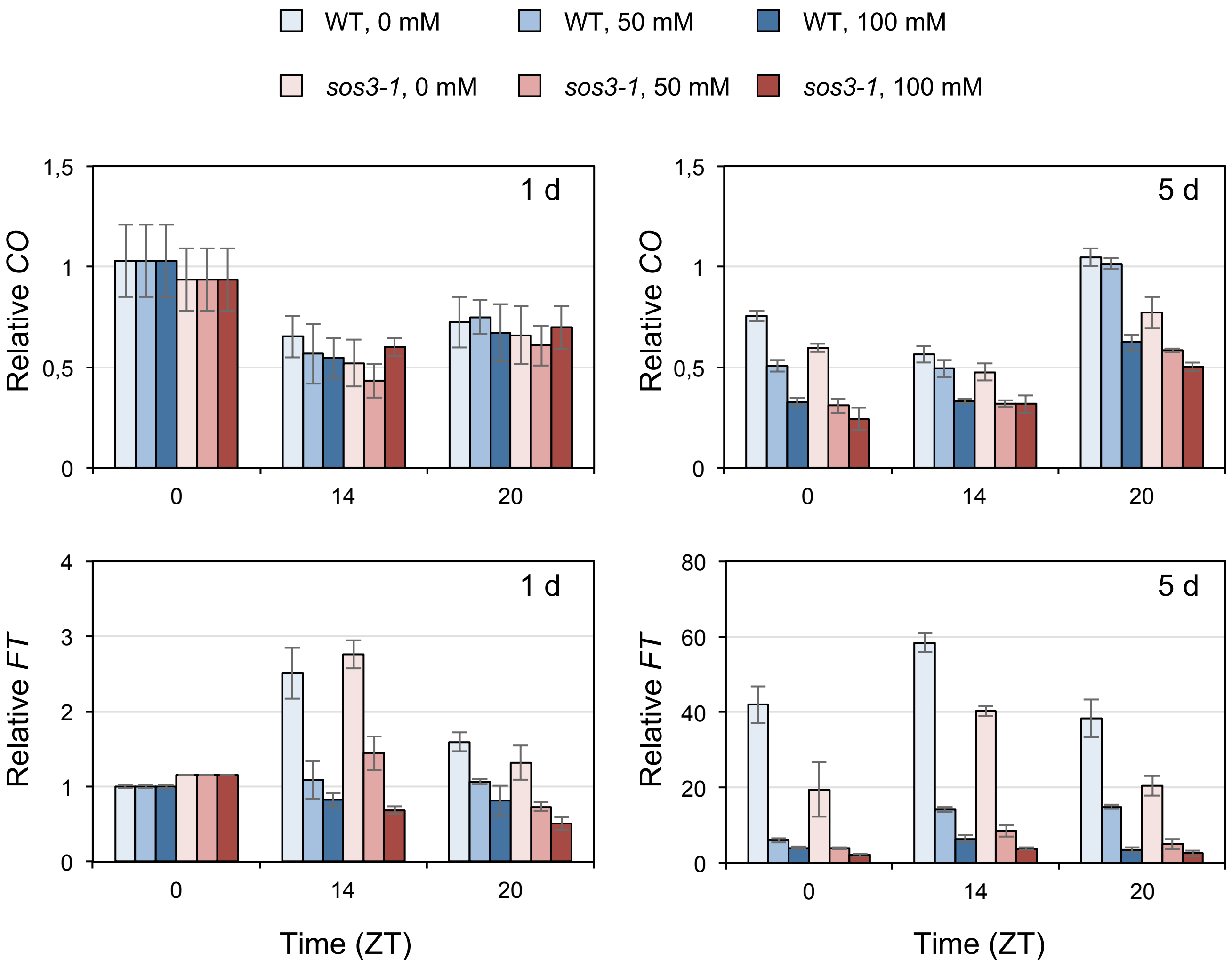


**Figure S11. Expression of *CO* and *FT* at 1- and 5-day under salt stress.**

Eight-day old plants of Col-*gl1* and *sos3-1* growing on ½ MS media (1% sucrose) under long-day conditions were treated with 100 mM NaCl at ZT0, and harvested at ZT14 and ZT20 of the same day (1 d), and at ZT0, ZT14, and ZT20 of the 5$^{th}$ day (5 d) after salt treatment. Transcript levels of *CO* and *FT* were measured by qRT-PCR and normalized to that of *At5g12240*. Error bars represent the SEM from three biological replicates, each with three technical replicates.
